# Immunothrombolytic monocyte-neutrophil axes dominate the single-cell landscape of human thrombosis

**DOI:** 10.1101/2024.01.10.574518

**Authors:** Kami Pekayvaz, Markus Joppich, Sophia Brambs, Viktoria Knottenberg, Luke Eivers, Alejandro Martinez-Navarro, Rainer Kaiser, Nina Meißner, Badr Kilani, Sven Stockhausen, Aleksandar Janjic, Vivien Polewka, Franziska Wendler, Augustin Droste zu Senden, Alexander Leunig, Michael Voelkl, Bernd Engelmann, Moritz R Hernandez Petzsche, Tobias Boeckh-Behrens, Thomas Liebig, Martin Dichgans, Wolfgang Enard, Ralf Zimmer, Steffen Tiedt, Steffen Massberg, Leo Nicolai, Konstantin Stark

## Abstract

Thrombotic diseases remain the major cause of death and disability worldwide with insufficient preventive and therapeutic strategies available. In the last decades a prominent inflammatory component has been identified as a key driver in the initiation and propagation of thrombosis – named thromboinflammation. However, a comprehensive investigation of the human immune system in thromboinflammation, beyond histological quantification, is lacking, which is essential for the development of novel therapeutic approaches. We therefore mapped the trajectories, functional states, and intercommunication of immune cells in stroke thrombi, retrieved by thrombectomy, at single-cell resolution. We reveal distinct leukocyte subpopulations with prothrombotic and, surprisingly, prominent fibrinolytic properties characterized by aberrant activation of intracellular host defense as well as hypoxia induced pathways. A prominent thrombolytic PLAU^high^, PLAUR^high^, THBD^high^ thrombus neutrophil subset, also expressing high levels of pro-recanalizing VEGFA and VEGFB, dominated the thrombus neutrophil environment. On the other hand CD16^high^ NR4A1^high^ non-classical monocytes with strong CXCL8, CXCL2, CXCL1 and CXCL16 mediated neutrophil-attracting and PLAU, PLAUR, THBD and TFPI mediated thrombolytic properties defined the thrombus monocyte environment. These thrombus monocyte subsets were characterized by high expression of TIMP1 and TREM1. These novel innate immune-cell subsets provide insights into the thrombogenic and pro-resolving properties of innate immune-cells. To provide mechanistic insight into these multi-omic findings, we utilized reverse translation approaches. *In vitro* as well as murine *in vivo* thrombosis models underlined the causal relevance of these immune-cell axes for thrombolysis: NR4A1^high^ thrombus monocytes acquired a neutrophil-chemoattractive transcriptomic phenotype, neutrophils continuously infiltrated established murine thrombi *in vivo* and acquired a HIF1α-mediated thrombolytic phenotype *in vitro*. A depletion of NR4A1^high^ thrombus monocytes reduced thrombus neutrophil influx and exacerbated thrombosis *in vivo*.

Together, this unravels cross-communicating monocyte and neutrophil subsets with thrombus-resolving properties and provide a publicly accessible immune-landscape of thrombosis. This provides a valuable resource for future research on thrombo- inflammation and might pave the way for novel immune-modulatory approaches for prevention or resolution of thrombosis.

## Introduction

Thrombosis and thromboembolism are a major global cause of morbidity and mortality by causing myocardial infarction, stroke, and venous thromboembolism. Simultaneously, antithrombotic therapies carry a significant risk of bleeding complications and are only partially effective in preventing thrombotic events^1, 2^. A strong interdependence between inflammation and thrombosis has gathered increasing attention in recent years, a process termed ‘immunothrombosis’ (in physiological processes) or ‘thromboinflammation’ (in disease scenarios)^1^. Studies involving rodent thrombosis models identified multiple therapeutically targetable inflammatory mechanisms in both low- and high shear clot formation that set off a thrombotic *vicious cycle* between neutrophils, monocytes, platelets, and the coagulation system^3–8^. In humans, histological analyses confirm a high frequency of leukocytes, particularly neutrophils, in cerebral and coronary thrombi ^9–12^. While highly valuable for initiating the translation of the concept of thromboinflammation to humans, these studies are limited to descriptive morphological and numerical conclusions. Novel -omics approaches allow a more comprehensive analysis of the immune system on a single cell level and hence might provide a leap towards a better mechanistic understanding of immune cell driven diseases in humans^13–15^. We hypothesized that leukocyte subsets perform specialized tasks by activating host defense pathways and subsequently modulating their thrombotic environment. Therefore, we performed a state-of-the-art multi-omic study of human thrombi and mapped the trajectories, functional states, and intercommunication of immune cells in thrombosis at single-cell resolution. This approach allowed us to uncover the immune cell landscape of thromboembolism: We found heterogenous neutrophil and monocyte subsets with divergent activation patterns and functional properties. Neutrophil subsets were characterized by pattern recognition, complement, interferon and NETosis signatures. In parallel, we also found a prominent thrombolytic and angiogenic neutrophil population. Monocytes were characterized by an alternative phenotype with clot resolving features and neutrophil chemotactic properties. To dissect this unexpected pro-resolving monocyte-neutrophil axis mechanistically, we performed *in vitro* and *in vivo* experiments: These confirmed thrombus NR4A^high^ alternative monocytes to hold neutrophil chemoattracting and thrombus-resolving properties. In summary, the immune landscape of human thrombosis reveals novel thrombus-modulating innate immune cell axes and differential activation of host defense pathways.

## Results

### Study design

To uncover the inflammatory signature of human thrombosis, we analyzed thrombi and peripheral blood from stroke patients (Suppl. Table 1). This allowed standardized large-scale access to human thrombosis (from n=31 subjects) with rapid and standardized *ex vivo* processing protocols, particularly important for neutrophil assessment (see methods). For the comprehensive analysis of the immune landscape of thrombosis we performed multi-color flow cytometry to assess overall frequencies of immune cell subsets. For an unbiased approach, we used single cell RNA sequencing (scRNA-seq) and CITE-seq, in which we studied cell phenotypes, developmental trajectories, functional programs and cross-communication between immune subsets in thrombosis. We validated our key findings utilizing a bulk RNA-sequencing approach (prime-seq) of FACS sorted immune cells (prime-seq^16^) in independent subjects (Fig. 1a). In a reverse translational approach, we further performed an *in vivo* mouse thrombosis model and *in vitro* assays to uncover the underlying mechanisms behind the monocyte-neutrophil interplay, identified in human thrombi (Fig. 1a).

**Figure 1.**
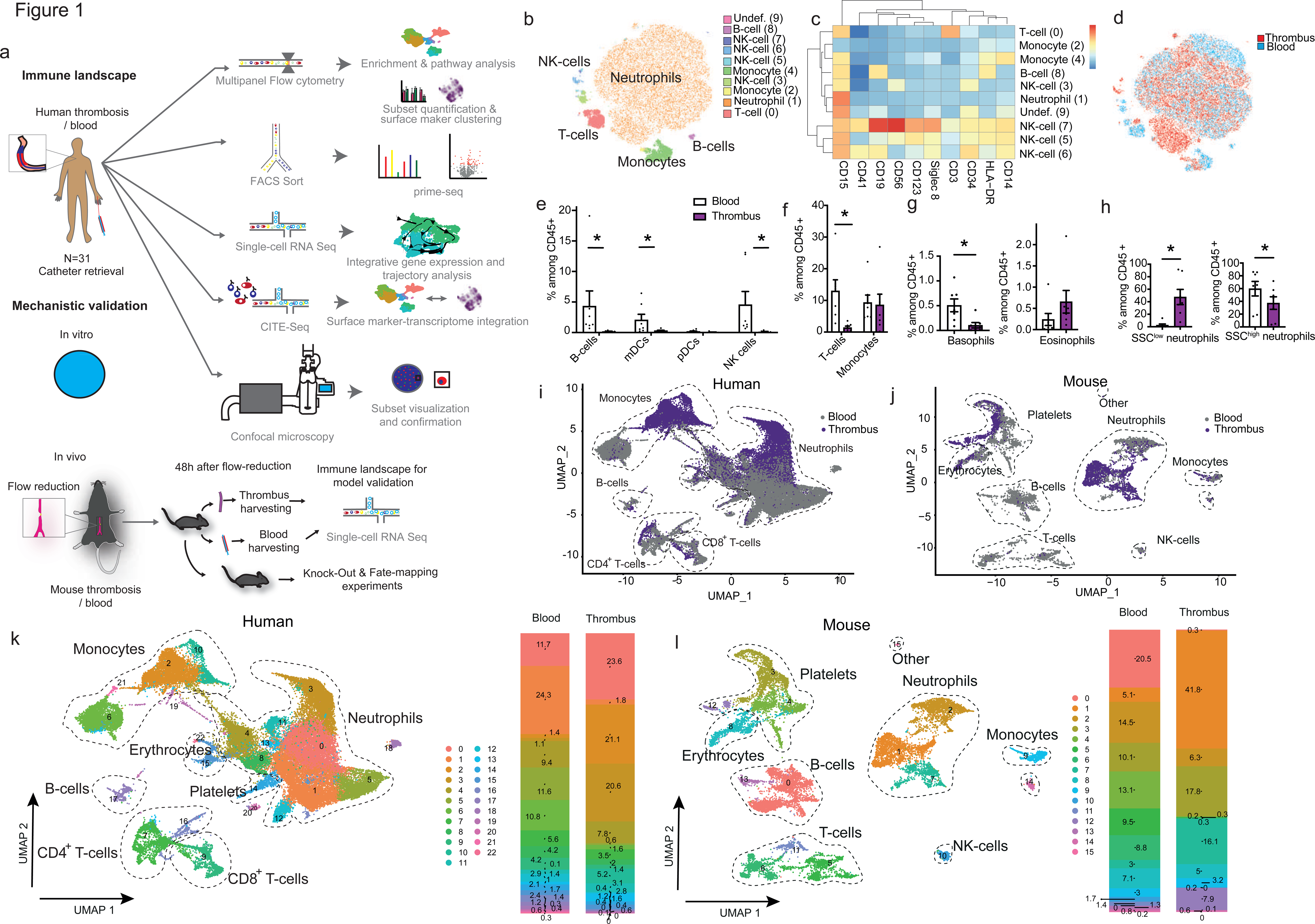
Thrombus immune-cell microenvironment (a) Study design: Human and mouse thrombosis was analysed by multiple -omic methods, including single-cell and CITE RNA-Seq, FACS-Sort with subsequent prime-seq^16^, multipanel flow-cytometry as well as confocal imaging. In a reverse translational approach, key pathways identified were subsequently confirmed by *in vitro* and *in vivo* approaches. Samples from a total of N=31 patients were included into the study. **(b)** Flow cytometry based immune-cell phenotyping of human thrombosis. t-SNE dimensionality reduction, cluster-identification with FlowSOM. (**c**) Heat map of the surface marker expression of each subcluster relative to the maximum expression of the surface marker. See Methods for the exact clustering of the heat map. **(d)** Colour coded t-SNE, dividing cells by origin from either thrombus or blood. **(b)-(d)** Depicts one representative thrombus and matched blood. **(e)**-**(h)** Quantification of immune-cells relative to all CD45^+^ leukocytes in thrombus compared to blood. Flow-cytometry phenotyping. n=7 thrombi & blood. Paired t-test was used for normally distributed data, Wilcoxon matched-pairs signed rank test was used for non-parametric distributed data. *P<0.05. **(i)**, **(j)** UMAP of human and mouse blood and thrombus scRNA-seq data coloured by origin (purple depicts thrombus-derived cells, grey depicts blood-derived cells). **(k)**, **(l)** UMAPs of human and mouse thrombus and blood scRNA-seq data coloured by cell clusters and cluster frequencies as separated by origin from blood or thrombus. **(i)**-**(l)** scRNA-seq analyses. Human UMAP based on: n=7 thrombi, n=6 blood. Mouse UMAP based on: n=3 thrombi, n=3 blood from respective mice (matched), n=3 blood from mice without thrombi despite flow-reduction (the latter cells depicted in Suppl. Fig. 3d).

### Single cell-based transcriptomics and surface profiling reveal prominent shifts between blood and thrombus with specialized immune-subsets conserved across species

First, to determine the leukocyte subsets involved in thrombosis, we utilized a flow-cytometry based phenotyping approach with subsequent dimensionality reduction by tSNE and FlowSOM based unsupervised clustering (Fig. 1b, c, Suppl. Fig. 1, 2). Paired comparisons of thrombi with blood from the same patients revealed distinct quantitative differences between blood and thrombi (Fig. 1d-h, Suppl. Fig. 1, 2): We found relative enrichment of distinct neutrophil subsets in thrombi, while most other leukocyte populations, including CD3^+^ T-cells, CD19^+^ B-cells, or CD56^+^ NK-cells and myeloid dendritic cells were less frequent in thrombi compared to blood. CD123^+^ basophils but not eosinophils were depleted in thrombi compared to blood, while plasmacytoid dendritic cell frequencies and monocyte counts stayed constant between blood and thrombi. Thrombus neutrophils showed a distinct sidewards scatter (SSC)^low^ phenotype (Fig. 1d-h).

To generate a comprehensive immune cells atlas and further delineate these phenotypic differences, we performed scRNA seq of retrieved thrombi and simultaneously of collected peripheral blood leukocytes. UMAP based dimensionality reduction showed strong shifts in the immune cell phenotype (Fig. 1i). ScRNA-seq analysis of a mouse model of thrombosis, induced by blood flow-reduction confirmed these strong phenotypic shifts compared to circulating innate immune cells (Fig. 1j), mimicking low-flow thrombus formation in the left atrial appendage during atrial fibrillation, the major driver of stroke in our patient cohort (Suppl. Table 1)^1, 3, 4^. Further elaboration on immune cell subclusters comparing blood and thrombi confirmed the predominance of innate immune cells, which also underwent the strongest changes in human and mouse thrombosis and in high shear as well as in low shear human thrombosis (Fig. 1k, l, Suppl. Fig. 3a-h).

### Thrombus neutrophils possess distinct thromboinflammatory and pro-resolving properties

Neutrophil-specific surface marker profiling recapitulated a CD15^+^ SSC^low^ neutrophil subset, that was significantly enriched in thrombi and further defined by lower CD16 expression in comparison to SSC^high^ neutrophils (Suppl. Fig. 4a, b). Unsupervised, fine-grained subclustering of the thrombus neutrophil (tNeutro) landscape by scRNA seq revealed more detailed insights into thrombus and blood neutrophil heterogeneity (Fig 2a-c). In total, we identified 13 neutrophil subclusters (Fig 2a-c, Suppl. Fig. 4c). Particularly VEGFA^high^ cluster 2 was confined to thrombi. Simultaneously, SSRGN^high^ Cluster 0, FN1^high^ ; NEAT1^high^ cluster 7 and ribosomal protein (RPL)^high^ cluster 8 were enriched in thrombi, whereas MX1^high^ Cluster 3 and mitochondrial gene (MT)^high^ cluster 6 showed comparable frequency in blood and thrombi (Fig. 2a-c, Suppl. Fig. 4c). Clusters 0,2,3,7 and 8 accounted for ∼ 90% of profiled thrombus Neutrophils (tNeutros). To allow an integrative analysis of surface proteome and transcriptome of neutrophils, we performed Cellular Indexing of Transcriptomes and Epitopes by Sequencing (CITE-Seq)^17^ in a subset of thrombi. This confirmed correlation of neutrophil clusters with low CD16 surface expression with low CD16 (*Fcgr3a*) gene expression, connecting transcriptomic with proteomic analyses (Suppl. Fig. 4d).

**Figure 2.**
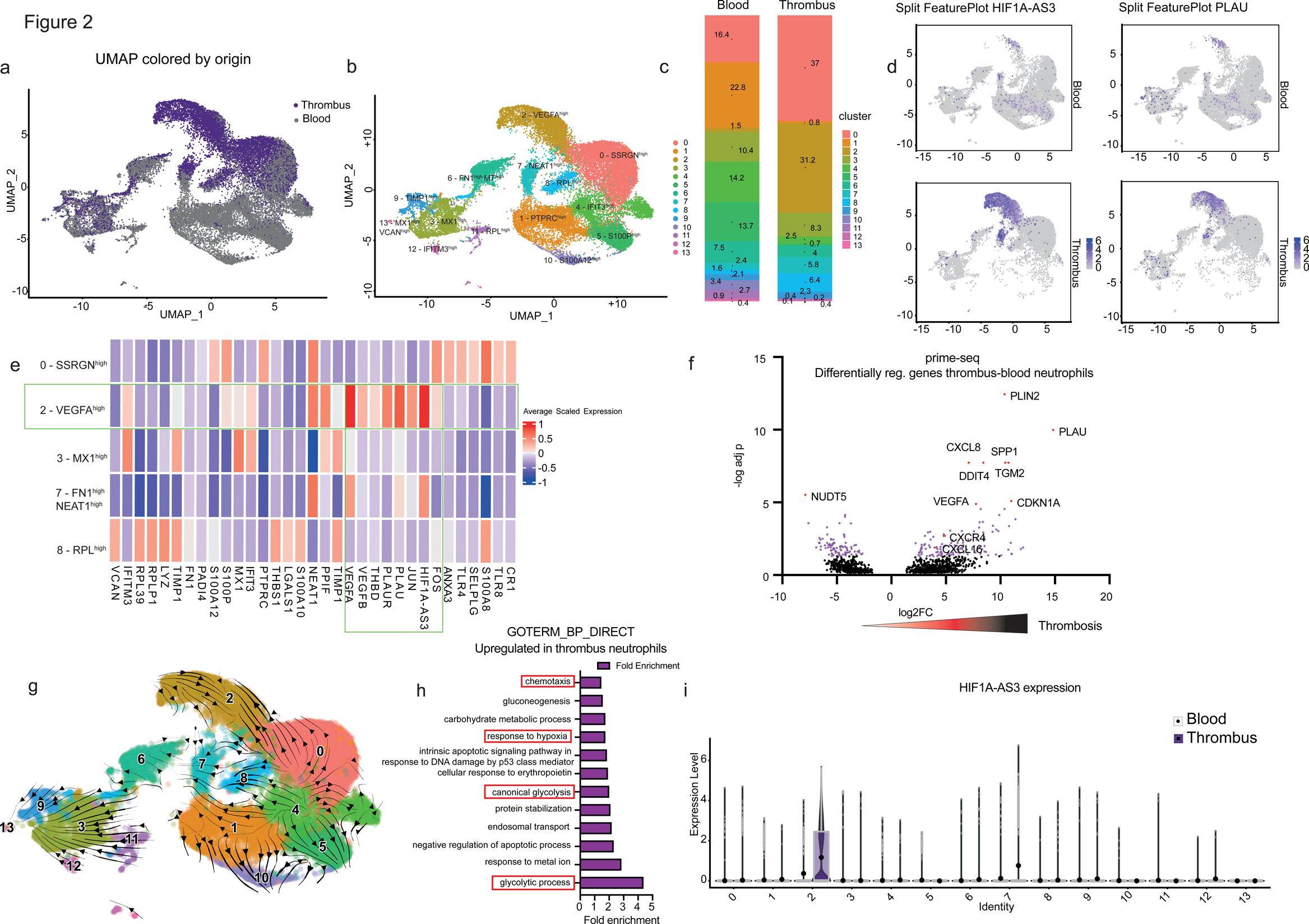
A detailed neutrophil atlas delineates subsets with prominent thrombus resolving and subsets with prothrombotic properties. (a) UMAP of human blood and thrombus neutrophils, reclustered from Figure 1k thrombi, coloured by origin (purple depicts thrombus-derived neutrophils, grey depicts blood-derived neutrophils). **(b)** UMAP of human thrombus and blood single-neutrophils coloured by neutrophil clusters. **(c)** Frequencies of neutrophil clusters in blood and thrombus in %. **(d)** FeaturePlots of HIF1A-AS3 and PLAU expression in blood and thrombus neutrophils. **(e)** Heatmap depicting selected cluster defining genes with functional relevance of cells from thrombus clusters or joint blood-thrombus clusters. Cluster 2, the main thrombus neutrophil cluster, and its respective functional cluster defining genes, is highlighted by a green box. **(f)** Volcano plot of differentially regulated genes between blood and thrombus neutrophils as determined by prime-seq^16^ of FACS-Sorted neutrophils. **(g)** Velocity analysis by scVelo. Arrows are depicting the calculated velocity vector field displayed as streamlines to determine cellular fate trajectories. **(h)** Goterm_BP_Direct analysis with genes enriched in thrombus neutrophils compared to blood neutrophils. n=8 thrombi, n=9 blood. **(i)** HIF1-AS3 expression across different neutrophil subclusters as determined by single-cell RNA Seq, subclusters are further depicted in Fig. 2. prime-seq^16^ analyses of FACS-sorted neutrophils: n=8 thrombi, n=9 blood (independent cohort).

Next, we sought to analyze the dominant tNeutro clusters in detail. We found that tNeutros expressed high levels of hypoxia associated genes such as *Hif1a-As3* in addition to genes associated with a thrombus-resolving phenotype such as *Plau* (Fig. 2d). SSRGN^high^ Cluster 0 was enriched in inflammatory transcripts like TLR4 and 8, as well as S100A proteins and receptors like complement receptor 1 and P-selectin ligand (Fig. 2e, Suppl. Fig. 4c). Many of these enriched genes are also implicated in pro-thrombotic NETosis^18–22^. MX1^high^ Cluster 3 showed upregulation of interferon stimulated genes (ISGs) like MX1, IFIT3 and IFITM3, also pointing towards a host defense signature. RPL^high^ Cluster 8 abundantly expressed ribosomal genes, indicating high translational activity, as well as pro-thrombotic thrombospondin (THBS1) transcription (Fig. 2e, Suppl. Fig. 4c).

*Fn1* (encoding fibronectin), a glycoprotein centrally involved in hemostasis^23, 24^ and involved in clot stabilization when incorporated into the fibrin network^24^ was expressed predominantly by FN1^high^ cluster 6 (Suppl. Fig. 5a). The transcriptome of cluster 6 was dominated by high FN1 expression. S100P^high^ cluster 5 and S100A12^high^ cluster 10 also expressed high levels of *Padi4*, likely representing pro-NETotic neutrophils^25–30^, fueling thrombosis^1^ (Suppl. Fig. 5a). This indicates the presence of multiple neutrophil clusters, which upregulate proteins known to drive thromboinflammation and propagate thrombosis.

Interestingly this contrasted with thrombus-exclusive VEGFA/B^high^ cluster 2, which was characterized by high expression of transcription and differentiation factor *Fos* and by particularly high expression of hypoxia associated *Hif1a-As3,* as well as anti-thrombotic *urokinase (PLAU), urokinase receptor (PLAUR)* and *thrombomodulin (THBD)* (Fig. 2e). Feature plots confirmed high levels of hypoxia associated genes such as *Hif1a-As3* in addition to genes associated with a thrombus-resolving phenotype such as *Plau* in thrombi (Fig. 2d). We validated global transcriptomic changes of thrombus neutrophils by prime-seq^16^ of FACS- sorted neutrophils from thrombi and respective blood samples including prominent upregulation of pro-resolving pathways (*Plau, Vegfa*). Indeed, the most strongly enriched gene across thrombus neutrophils was *Plau* (the gene encoding urokinase) (Fig. 2f, Suppl. Fig. 5b). In summary, we define heterogenous populations of neutrophils in human thrombi, which differ significantly from circulating neutrophils. Moreover, our scRNA-seq data provide evidence of thrombus neutrophil subsets with diverse, potentially diametrical effector functions: Prothrombotic and NETotic populations as well as pro-resolving subsets with hypoxic attributes.

Due to these strong shifts culminating in separately clustering immune cell subsets, we asked whether neutrophils emerge from their phenotypically nearest blood counterparts. A cell trajectory inference by scVelo^31^ revealed anti-thrombotic thrombus neutrophils (VEGFA^high^ cluster 2) as well as other thrombus neutrophil subsets to emerge from common circulating blood neutrophils. The main vectors from possible blood progenitors, recovering dynamic information from splicing kinetics on a single cell level, were jointly directed towards the thrombus progeny as well as further blood neutrophil maturation stages (IFIT3^high^ cluster 4 and subsequently SSRGN^high^ cluster 0 developing towards VEGFA^high^ cluster 2) (Figure 2g). This suggests a continuous polarization of blood neutrophils towards thrombus neutrophils.

### Intrathrombotic hypoxia imprints a pro-resolving phenotype on continuously infiltrating blood neutrophils

To get more functional insight into global changes in thrombus neutrophils we performed a DAVID set enrichment analysis on the Gene Ontology biological process gene sets (GOTERM_BP_DIRECT). This revealed multiple hypoxia related pathways to be upregulated across thrombus neutrophils (Figure 2h). At single-cell resolution Hif1a-As3 was mainly confined to anti-thrombotic VEGFA^high^ cluster 2 (Figure 2i). We therefore hypothesized that the hypoxic environment within the thrombus might potentially trigger the distinct shifts in neutrophils towards a pro-resolving phenotype with thrombolytic potential. The expression of urokinase in thrombus neutrophils was indeed confirmed by immunofluorescence (Figure 3a). *In vitro*, we observed that pharmacological HIF-1α stabilization by Roxadustat recapitulated the shifts in the neutrophil compartment towards a CD16^low^ neutrophil subset (Figure 3b). These CD16^low^ neutrophils also had a SSC^low^ degranulated phenotype (figure 3c, d). To evaluate the transcriptionally observed strong fibrinolytic potential of hypoxic neutrophils, we analyzed the clearance of cross-linked fibrin coated surfaces. Indeed, neutrophils were able to continuously clear fibrin *in vitro* and this feature was significantly amplified by Roxadustat mediated hypoxia mimicking (Figure 3e, f). This indicates that thrombus hypoxia contributes to skewing tNeutros towards a thrombolytic phenotype.

**Figure 3.**
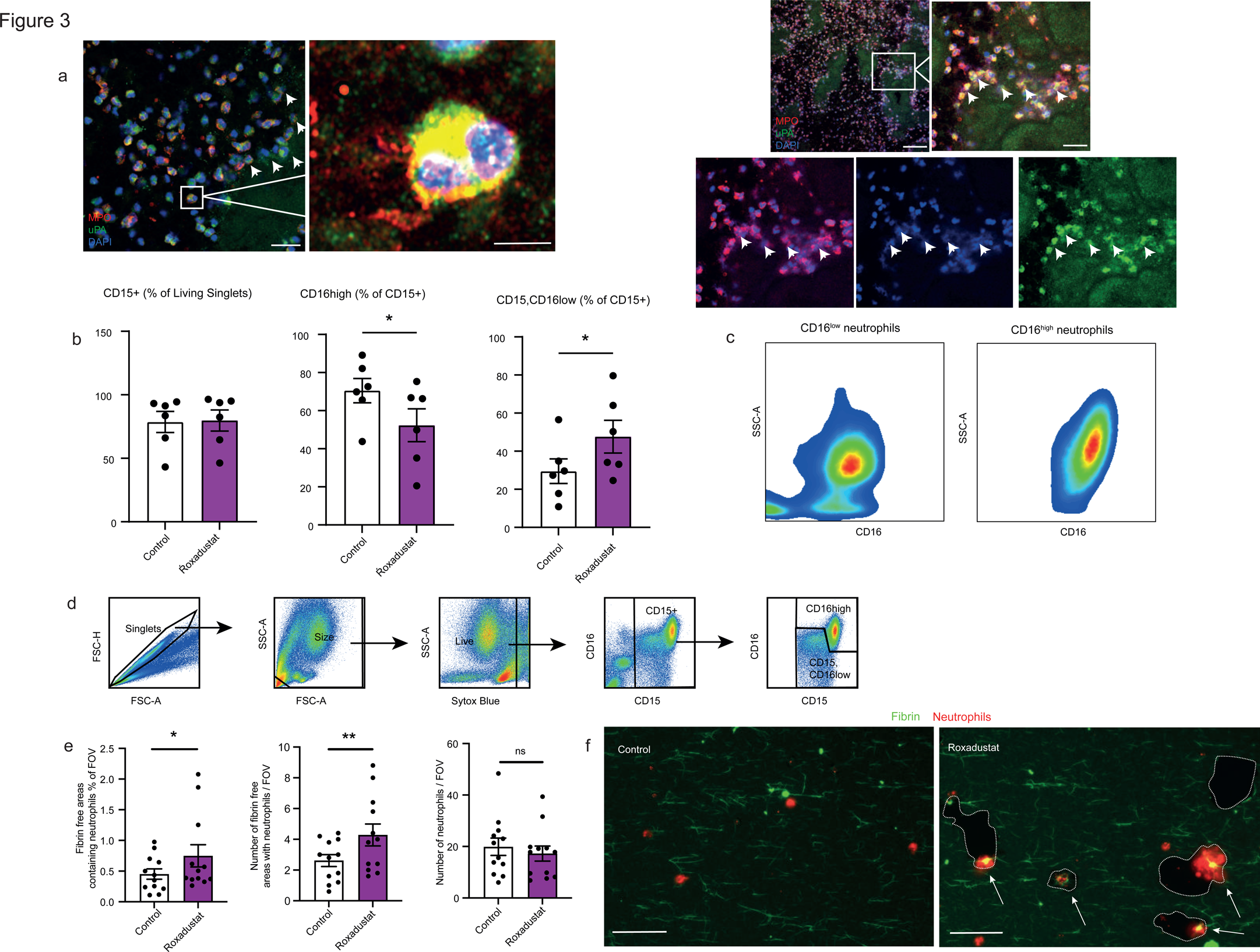
Hypoxia reproduces the CD16^low^ pro-resolving neutrophil phenotype (a) Immunofluorescence imaging of thrombus neutrophils (red) stained for urokinase (UPA) (green) and DNA (blue). Scalebar left: 25µm, scalebar middle: 5µm, scalebar right, overview: 100µm & scalebar right, magnified image: 25µm. **(b)** Analysis of CD16^high^ and CD16^low^ neutrophil counts (% of living singlets), comparing neutrophils after Roxadustat mediated HIF1α stabilization with control neutrophils. **(c)** Representative FSC-A and SSC-A gates of CD16^high^ and CD16^low^ neutrophils, representing the rather SSC^low^ phenotype of CD16^low^ neutrophils. n=6 independent experiments. **(d)** Flow-cytometry based gating strategy of neutrophil analysis after Roxadustat mediated HIF1α stabilization. **(e)** *In vitro* analysis of fibrinolytic potential of hypoxic neutrophils, triggered by pharmacological, Roxadustat mediated intracellular induction of HIF1α signaling. Analysis of fibrin free area, number of fibrin free areas and number of neutrophils. Paired t-test was used for normally distributed data, Wilcoxon matched-pairs signed rank test was used for non-parametric distributed data. n=12 replicates from n=4 independent experiments. **(f)** Representative images, depicting fibrin free areas with neutrophils, most prominently in Roxadustat treated wells. Scale bar: 50 µm. *P<0.05. ** P<0.01.

### Thrombus monocytes show a nonclassical-like phenotype with thrombus modulating properties and strong neutrophil-attracting features

Next, we focused on thrombus monocytes as they are the second most common leukocyte population in thrombi. Flow cytometry showed a shift in the monocyte compartment driven by increased CD16 surface expression (Figure 4a,b, Suppl. Fig. 6a). This points towards an alternative monocyte phenotype of thrombus monocytes. To gain insight into the global transcriptional landscape across tMono subsets, we performed bulk prime-seq^16^ on FACS/sorted monocytes from thrombi. We confirmed global upregulation of master transcription factor *Nr4a1*, as well as *Fcgr3a* (encoding CD16) in thrombus monocytes, jointly highlighting tMono differentiation along an alternative/non-classical trajectory (Fig. 4c). In addition, thrombus nonclassical like monocytes were characterized by upregulation of hypoxia/HIF-1α signaling, pro-resolving and fibrinolytic pathways, and most prominently an upregulation of neutrophil chemoattractants (Fig. 4c-e, Suppl. Fig. 6b, Suppl. Fig. 7a,b). Next, we dissected the thrombosis-related triggers of monocyte differentiation towards a tMono-like phenotype with thrombolytic and neutrophil-activating properties. Here, we focused on hypoxia and coagulation activation as potential drivers. Even though, mimicking hypoxia by Roxadustat did not recapitulate the shift in monocyte phenotype observed within the thrombi (Suppl. Fig. 7c), thrombin mediated coagulation of whole-blood *in vitro* recapitulated a shift of monocytes towards a phenotype with neutrophil-activating properties, indicating that the shift in monocyte phenotype might be directly dependent on the thrombus microenvironment (Fig. 4f).

**Figure 4.**
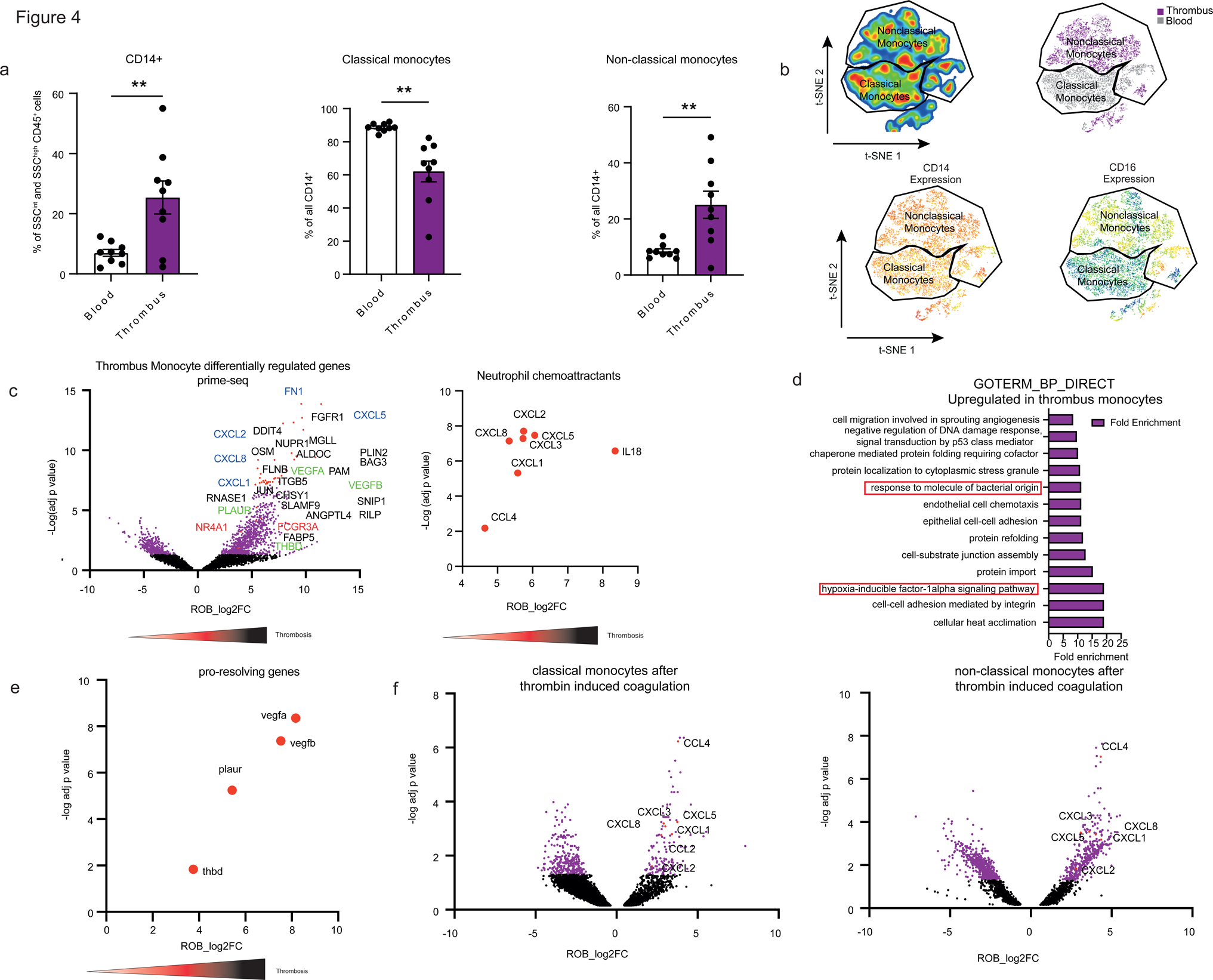
Thrombus monocytes collectively acquire a non-classical like, neutrophil chemoattracting phenotype. (a) Flow-cytometry based quantification of CD14^+^ cells among all CD45^+^ cells present in the ordinary SSC gates (SSC^int^ or SSC^high^) and further subdifferentiation classical and non-classical monocytes among all monocytes in blood and thrombi. n=9 blood and n=9 thrombi. **(b)** t-SNE based dimensionality reduction of monocyte flow-cytometry data further colour coded by origin (top right), CD14 expression (bottom left) and CD16 expression (bottom right). Paired t-test was used for normally distributed data. ** P<0.01. (**c**) Volcano Plot of differentially regulated genes between blood and thrombus monocytes as determined by prime-seq^16^ of FACS-sorted monocytes (left). Illustration of neutrophil active chemokines enriched in thrombus monocytes as determined by prime-seq^16^ of FACS-sorted monocytes (right). **(d)** Goterm_BP_Direct analysis with genes enriched in thrombus monocytes compared to blood monocytes. Red box depicts hypoxia-inducible factor- 1alpha signaling pathway and response to molecule of bacterial origin. **(e)** Volcano plot of prime-seq^16^ data of FACS/Sorted monocytes specifically depicting upregulated genes associated with recanalization / thrombolysis. **(c)-(e)** n=7 thrombi, n=7 blood. **(f)** Volcano plots of differentially regulated genes as determined by prime-seq^16^ of FACS-Sorted classical and nonclassical monocytes after thrombin and calcium mediated induction of the coagulation cascade. n=5/group.

To pinpoint monocyte heterogeneity at single cell resolution we performed single-cell RNA-seq and CITE-seq (Fig. 5a-c). Monocytes were clustered in 10 different subsets, with Clusters 0 (CLEC5A^high^), 3 (THBS1^high^), 4 (CXCR4^high^), 6 (CXCL8^high^), 7 (CSTD^high^) and 10 representing the most frequent thrombus monocyte (tMono) clusters (Fig. 5a-c, Suppl. Fig. 7d). Indeed, scVelo^31^ based differentiation vectors suggested a possible development of thrombus monocytes from respective blood progenitors. CD16 (FCGR3A)^high^ Cluster 9 emerged as the connecting link population between blood and thrombus monocytes as evidenced by the developmental trajectories (Fig. 5d). This trajectory was based on the expression and distribution of spliced versus unspliced variants of human genes such as *Eri1*, substantiating a development of blood monocytes towards the shared thrombus / blood cluster 9 monocytes (Fig. 5d, Suppl. Fig. 7e). Differential analysis revealed *Fcgr3a* (encoding CD16) to be strongly enriched beyond cluster 9 across major thrombus monocyte clusters (Fig 5e, Suppl. Fig. 7f). Along these lines, *Nr4a1*, the key transcription factor responsible for CD16^hi^CD14^low^ alternative monocyte differentiation^32^, was prominently expressed in thrombus monocytes (Fig. 5f).

**Figure 5.**
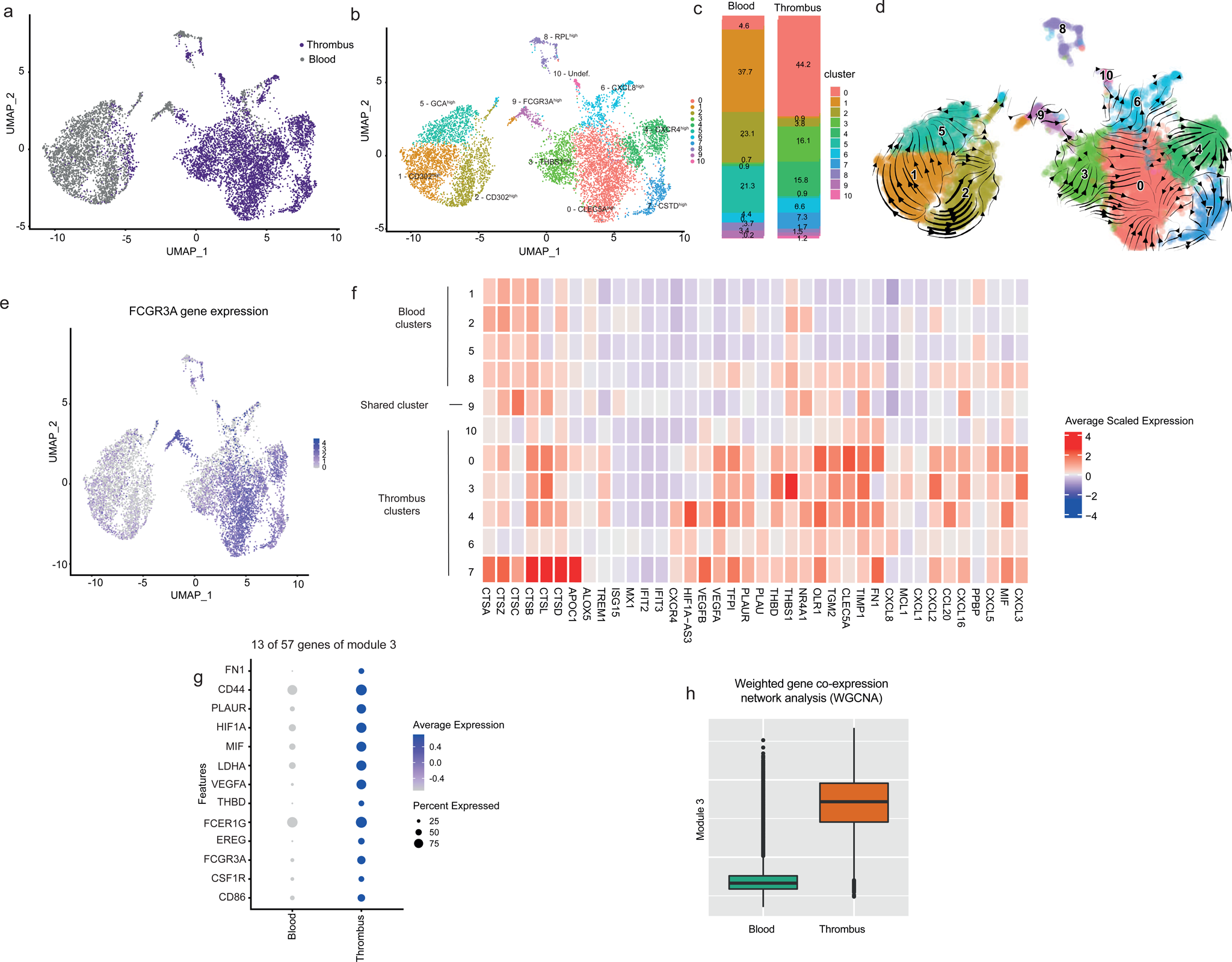
Single-cell resolution delineates multiple functional monocyte clusters with prothrombotic, thrombolytic and neutrophil attracting tasks that are present in human thrombi. (a) UMAP of human blood and thrombus monocytes, reclustered from Figure 1k, coloured by origin (purple depicts thrombus-derived neutrophils, grey depicts blood-derived neutrophils). **(b)** UMAP of human thrombus and blood single-monocytes with colored and annotated monocyte clusters. **(c)** Frequencies of monocyte clusters in blood and thrombus in %. **(d)** Velocity analysis by scVelo. Arrows are depicting the calculated velocity vector field displayed as streamlines to determine cellular fate trajectories **(e)** FeaturePlots of FCGR3A (CD16) expression in blood and thrombus neutrophils. **(f)** Heatmap depicting selected cluster defining genes with functional relevance of cells from thrombus, blood and joint blood- thrombus monocyte clusters. **(g)** Weighted Gene Co-expression Network Analysis (WGNCA) of monocytes, allowing identification of gene networks regulated in a joint manner across cells and tissues. Exemplary inclusion of 14 of a total of 57 genes within identified Module 3. Complete module is depicted in Suppl. Fig. 9. **(h)** Module Score of Gene Module 3 depicted across blood and thrombus, illustrated as box-plots showing median and interquartile range in blood and thrombus monocytes.

Next, we assessed thrombus monocyte phenotypes in more detail. All major thrombus monocyte subsets exhibited strong expression of neutrophil chemoattractants, i.e. *Cxcl8, Cxcl2, Cxcl5, Cxcl3,* and *MIF* (Fig. 5f). In addition, transcripts associated with thrombus resolution and recanalization: urokinase receptor, thrombomodulin, vascular endothelial growth factor A and B and tissue factor pathway inhibitor (*Plaur*, *Thbd*, *Vegfa*, *Vegfb*, *Tfpi*), were highly expressed across all major thrombus monocyte subsets, most prominently by CXCR4^high^ cluster 4, as well as THBS1^high^ cluster 3 and CLEC5A^high^ cluster 0 thrombus monocytes (Fig. 5f, Suppl. Fig. 7d). Also, thrombus monocytes, particularly CXCR4^high^ Cluster 4, expressed high levels of Hif1-As3 indicating hypoxia-induced pathway induction as observed in tNeutros. Multiple cathepsins were enriched in CTSD^high^ cluster 7, suggesting a role of this cluster in extracellular matrix degradation as well as lysosomal protein digestion (Fig. 5f). Interestingly, the CTSD^high^ cluster 7 also showed prominent upregulation of APOC1, which is activated when monocytes transdifferentiate to macrophages^33^ (Fig. 5f). In summary, thrombus monocytes exhibited distinct phenotypes, while sharing (1) an alternative monocyte phenotype, (2) clot resolving features and (3) expression of neutrophil attracting factors. Next, we asked if these features were indeed jointly regulated across monocytes in the sense of directed cellular programs. To this end, we employed weighted gene correlation network analysis (WGCNA) to distill coordinative cellular responses as condensed “gene modules”^34, 35^. WGCNA analysis revealed a total of 8 modules of varying size (Fig. 5g, Suppl. Fig. 8).

Modules 3-6 were enriched in monocytes within the thrombus (Fig. 5h, Suppl. Fig. 9) with module 3 depicting the most prominent enrichment within thrombi. Interestingly, module 3 consisted of neutrophil chemoattracting (*Mif*), thrombus resolving (*Plaur, Thbd*) as well as alternative monocyte defining genes (*Fcgr3a*). This indicates a possible joint upstream regulation of the phenotypic shift in thrombus monocytes (Fig. 5g, h).

In summary, thrombus monocytes are defined by distinct clusters mainly showing a non-classical like phenotype with parallel upregulation of neutrophil chemotactic signals and thrombus modulating properties.

### Identification of a pro-resolving monocyte-neutrophil axis in thrombosis

To allow a detailed understanding of the intercellular immune cell communication within thrombi, a single-cell interactome analysis was performed. In line with the strong expression of neutrophil active chemokines by monocytes, the chemokine interactome outlined abundant intercellular communication pathways between monocyte and neutrophil subsets, strongly mediated by the CXCL8-CXCR1 / CXCR2 axis (Fig. 6a, Suppl. Fig. 10).

**Figure 6.**
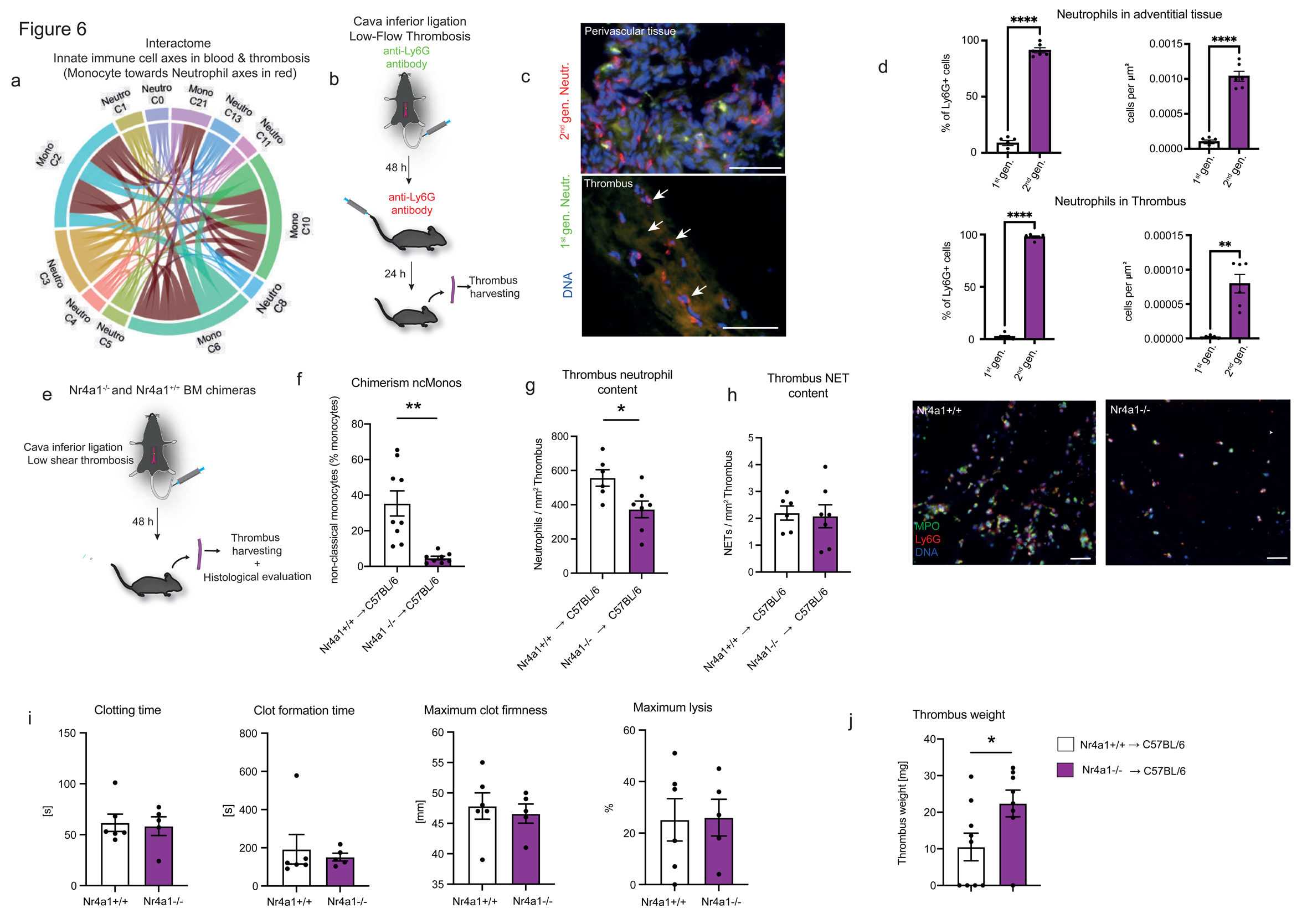
Reverse translation confirms non-classical like thrombus monocytes to fuel continuous blood neutrophil recruitment, resolving thrombosis. (a) Chemokine interactome between human neutrophils and monocytes analysed for clusters further specified in Figure 1 & Suppl. Fig. 3. The connections in dark red highlight monocyte -> neutrophile interactions. **(b)** Murine model of thrombosis. Illustration depicting the experimental setup: FITC tagged Anti-Ly6G neutrophil antibody was injected during inferior vena cava ligation. 48h after inferior vena cava ligation PE tagged Anti-Ly6G antibody was injected. 24h after the second antibody injection, mice were sacrificed and thrombi harvested and analyzed histologically. **(c)** Representative epifluorescence images of 1^st^ generation neutrophils (green) and 2^nd^ generation neutrophils (red) and DNA (blue) within the thrombus and within the perivascular tissue. Scalebar: 50 µm. **(d)** Quantification of 1^st^ and 2^nd^ generation neutrophils in % of all neutrophils and as cells/µm^2^ in the vessel wall (top) and within the thrombus (bottom). n=6 mice. **(e)** Illustration depicting the experimental setup: Cava inferior ligation was performed to induce thrombosis by flow-reduction. 48h after flow-reduction, chimeric mice were sacrificed, thrombi were harvested and histologically analysed. **(f)** Analysis of chimerism in Nr4a1^-/-^ mice by determining LY6C^low^ non-classical monocyte counts in blood by flow-cytometry. n=8-9 mice **(g)** Thrombus neutrophil counts per mm^2^ in Nr4a1^-/-^ and Nr4a1^+/+^ bone marrow chimeras. n=6-7 mice. **(h)** Thrombus NET content per mm^2^. n=6-7 mice. Right: Representative epifluorescence image of Ly6G^+^ MPO^+^ neutrophils in Nr4a1^-/-^ and Nr4a1^+/+^ bone marrow chimera thrombi. Scalebar: 50µm **(i)** ROTEM analysis of blood from Nr4a1^-/-^ and Nr4a1^+/+^ mice: Clotting time (left), clot formation time (left middle), maximum clot firmness (right middle), maximum lysis (right). n=5-6 mice. **(j)** Thrombus weight analysis of Nr4a1^-/-^ and Nr4a1^+/+^ bone marrow chimera thrombi in mg. n=8-9 mice. **(d)** Paired t-test was used. **(f) – (j)** Unpaired t-test was used for normally distributed data, Mann-Whitney test was used for non-parametric distributed data. *P<0.05. ** P<0.01.

Due to this prominent neutrophil-chemotactic phenotype of tMonos and the presence of heterogenous neutrophil populations within thrombi, we hypothesized that there might be a secondary, tMono driven influx of blood neutrophils during later stages of thrombosis.

To investigate this hypothesis further, we opted to utilize the established murine inferior vena cava stenosis model (Fig. 1)^4^ mimicking low-flow thrombosis found in human cardioembolic stroke. We focused on low-flow thrombosis as most of the investigated human thrombi were of cardioembolic origin (compare Suppl. Table 1). In this murine thrombosis model, monocytes underwent similar phenotypic changes as their human counterparts: Again, several neutrophil chemoattractants, including *Cxcl2*, *Ccl2*, *Mif*, *Cxcl16* and the cytokine *Il1b* were enriched in thrombus monocytes (Suppl. Fig. 11). A prominent *Ccr2^high^*subset (Cluster 9) and a smaller *Cx3cr1^high^* subset (Cluster 14) were present in blood as well as in thrombus (Suppl. Fig. 11b). *Ly6c2* was strongly downregulated in the major thrombus monocyte subset compared to its blood counterpart (Cluster 9). This corresponded to the phenotypic shift towards a non-classical like CD16 (*Fcgr3a*)^high^ phenotype observed in human thrombi (Suppl. Fig. 11a, b)^36^, indicating that the murine disease model mimics the human counterpart.

Next, we investigated possible continuous neutrophil recruitment towards developing thrombi. Therefore, we employed a neutrophil fate mapping approach in our flow reduction induced murine thrombosis model, allowing investigation of the temporal dynamics of neutrophil recruitment. We made use of a multi-color labelling approach – the first surge of neutrophils was labelled with a green fluorescent antibody (1^st^ generation neutrophils). 48h after the first labelling, neutrophils were labelled again with a red fluorescent dye (2^nd^ generation neutrophils), allowing differentiation between first and second surge of neutrophils into the thrombus at 72h after thrombus induction (sacrifice and organ harvesting) (Fig. 6b). Within the perivascular tissue, a green and a red fluorescing neutrophil population was detectable, indicating a successful fate mapping approach (Fig. 6c). In thrombi, 1^st^ gen. neutrophils were rarely detectable, but 2^nd^ gen. neutrophils were present at high frequencies. This indicates that neutrophils infiltrate the thrombus also at later stages, continuously replacing short-lived earlier thrombus neutrophil populations, which might no longer be detectable due to NETosis^1^ (Fig. 6d). Next, based on the human and mouse scRNA-seq data we hypothesized that human CD16 (*Fcgr3a*)^high^ monocytes, corresponding to murine LY6C^low^ monocytes, substantially fuel this continuous thrombus neutrophil influx by their above-mentioned prominent expression of neutrophil chemoattractants.

The transcription factor *Nr4a1* essentially mediates the transition towards a LY6C^low^ monocyte phenotype^32^, thereby providing a tool for genetic depletion of non-classical thrombus monocytes. We used *Nr4a1^-/-^* bone marrow chimeras to assess contribution of thrombus monocytes to neutrophil accumulation within thrombi (Fig. 6e). In line, genetic deletion of Nr4a1 in bone marrow derived cells depletes nonclassical monocytes as described^37^. Flow- cytometric analysis confirmed a depletion of circulating LY6C*^low^* monocytes (Fig. 6f, Suppl. Fig. 12). Analysis of low-flow thrombi indeed revealed a strong reduction in neutrophil content in the chimeric *Nr4a1^-/-^* thrombi compared to *Nr4a1^+/+^* chimeras, confirming non-classical like monocytes driven continuous influx of blood neutrophils into thrombi (Fig. 6g). Thrombus NET content remained unchanged (Fig. 6h). Assessment of clotting or clot formation time, maximum clot firmness and maximum lysis by thrombelastometry excluded a significant direct difference in coagulation between *Nr4a1^-/-^*chimeras and WT mice *in vitro.* This underlines possible indirect effects of non-classical like monocytes by recruiting neutrophils *in vivo during the thrombus resolution phase* (Fig. 6i). Indeed, *in vivo*, the lack of secondary neutrophil infiltration into thrombi in *Nr4a1^-/-^* chimeras resulted in increased thrombus weight (Fig. 6″j).

In summary, this indicates an unexpected pro-resolving monocyte-neutrophil axis fueled by *Nr4a1^high^*tMonocytes continuously recruiting a secondary, blood-borne neutrophil population into the thrombus which become hypoxic and subsequently polarize into thrombolytic and angiogenic tNeutros.

## DISCUSSION

The contribution of immune cells to thrombosis has become a center of attention in multiple diseases beyond cardiovascular disease, most importantly in COVID-19, but also in autoimmunity, sepsis and cancer^1, 38, 39^. The contribution of immune cells to thrombosis has been extensively studied in animal models in the past decades, deeming innate immune cells strongly prothrombotic at early phases of thrombogenesis^1,4^. However, translational approaches that allow a comprehensive cellular and functional understanding of the immune landscape in human thrombosis have been lacking. The exact leukocyte-phenotypes during thrombotic transition in respect to classical blood phenotypes remained unclear. Functionally heterogeneous neutrophils and monocytes are the most enriched leukocyte subsets within human thrombi. Simultaneously neutrophils and monocytes - unlike NK-cells, T-cells or B-cells

- undergo strong phenotypic shifts in thrombi compared to their blood counterparts as evident by the transcriptome and surfaceome. Velocity analysis suggests that these cells develop from distinct blood progenitors. We show that intercommunicating neutrophil and monocyte subsets hold highly specialized thromboinflammatory, and, surprisingly, immunothrombolytic features in humans and in mice. Concretely, thrombus monocytes jointly develop from a common non-classical blood monocyte progenitor and form a heterogeneous thrombus monocyte landscape, which is however uniformly characterized by its CD16^high^ NR4A1^high^ phenotype with strong neutrophil chemoattracting features across all thrombus monocyte subsets. Simultaneously, thrombus monocytes, in contrast to circulating blood monocytes, show great thrombus-modulating properties, with different cluster specific functions. Monocyte cluster 3 holds strong thrombolytic and pro-resolving properties with high *Thbd*, *Plaur* and *Thbs1* expression. Whereas thrombus monocyte cluster 7 shows high expression of FN1 and cathepsins, both associated with thrombus formation^23, 24, 40, 41^. A weighted gene co-expression network analysis suggests these monocyte programs to be regulated in a joint, upstream manner across single-cells. However, coagulation activation, but not hypoxia *per se*, recapitulates this non-classical monocyte phenotype *in vitro,* which points towards monocyte polarization driven by the thrombus microenvironment. This pro-resolving thrombus alternative monocyte phenotype is in line with previously described vasculoprotective properties of non- classical monocytes^42, 43^. In line with their strong neutrophil chemoattracting potential, non- classical NR4A1^high^ thrombus monocytes are characterized by high expression of the proinflammatory Triggering Receptor Expressed on Myeloid Cells (TREM1), a defining factor of immune-cell activation in host defence during sepsis^44^.

The human neutrophil landscape in thrombosis is highly diverse, with multiple prothrombotic subsets present in blood and within thrombi, enriched in pro-NETotic genes and with cellular states such as *Padi4* associated with an aberrant activation of host defense as seen in immunothrombosis^45^. On the other hand, a very prominent pro-resolving neutrophil subset is confined to thrombi and not present in the circulating neutrophil landscape (VEGFA^high^ Cluster 2 neutrophils). This VEGFA^high^ neutrophil subset is functionally reminiscent yet transcriptionally distinct of previously described proangiogenic neutrophils^46^. In addition to pro-resolving *Plaur*, *Plau* or *Vegfa, Vegfb*, the neutrophil cluster 2 is particularly enriched in hypoxia driven *Hif1a- As3*. Indeed, this distinct phenotype of neutrophil cluster 2 can be reproduced by pharmacological hypoxia induction. In earlier mouse studies, HIF1α correlated negatively with oxygen tension in the thrombus, but positively with *Vegfa* expression. *Vegfa* is also induced upon HIF1α stimulation, supporting thrombus resolution and recanalization^47, 48^. Augmenting local physiological immune-thrombolytic pathways might hence facilitate effective thrombolysis without triggering major systemic side effects. Indeed, Roxadustat driven induction of intracellular hypoxia pathways not only triggers a phenocopy of CD16^low^ thrombus neutrophils by flow-cytometry but also recapitulates an immune-thrombolytic neutrophil behaviour *in vitro*.

Mimicking human low-flow thrombosis, a flow restriction thrombosis model in mice recapitulates human immune cell composition and phenotype on a single-cell level. This allows mechanistic anaylsis and manipulation of the observed monocyte-neutrophil cross-communication. Since thrombus monocytes hold strong neutrophil chemoattracting properties, we hypothesized that blood neutrophils might continuously infiltrate the thrombus.

A fate-mapping approach shows neutrophils to continuously infiltrate established thrombi. Depletion of NR4A1^high^ neutrophil chemoattractive thrombus monocytes significantly reduces this thrombus neutrophil influx. Further, the depletion of NR4A1^high^ monocytes exacerbates thrombosis. This suits the neutrophil attracting properties described for *Nr4a1* dependent LY6C^low^ monocytes^32^ and their contribution to vascular homeostasis under physiological conditions: Scavenging endothelial debris in the vasculature^49^. The chemokine mediated recruitment of pro-resolving neutrophils further confirms early reports on thrombus resolution upon *in vivo* interleukin-8 administration^50^ and an impaired thrombus resolution during neutropenia^51^.

VEGFA^high^ neutrophils strongly upregulate *Plau* encoding the gene product urokinase within the thrombotic environment, which is a plasminogen activator. Urokinase as well as other plasminogen activators are most efficiently being used by neurologists to treat strokes, reduce thrombus burden and rapidly restore cerebral blood to significantly reduce death and disability however these are accompanied by a systemic bleeding risk with possible dire consequences^52, 53^. An intrinsic thrombolytic phenotype of distinct neutrophils, characterized by upregulation of these classical hospital-provided therapeutic agents, is particularly intriguing as it outlines intrinsic thrombus-resolving properties by leukocytes trapped within thrombi. Therefore, this identifies the therapeutic manipulation of immune-cells, phenocopying distinct thrombolytic immune-phenotypes, as a novel option to support local thrombolysis without risking systemic bleeding side-effects.

In summary, this maps the immunological landscape of thrombo-inflammation, provides a resource for future research in vascular immunology and benchmarks novel, highly specialized immune-cell phenotypes that underline the plasticity of immune-cells in resolving vascular disease. Beyond the descriptional multi-omics approach the presence of a pro-resolving NR4A1-driven monocyte axis in murine thrombosis -closely resembling its human counterpart- confirms previous descriptional multi-omic observations.

This provides novel, so far unknown neutrophil and monocyte phenotypes which hold important ramifications for immune-cell plasticity, acquiring protective, therapeutically exploitable roles in the context of resolving deadly vascular events. Further, this dataset holds important directions for future research on the immunologic aspect of thrombosis outlining multiple novel thrombolytic but also pro-thrombotic immune-cell phenotypes as possible targets that allow circumvention of the increased bleeding risks in conventional therapies^1, 2, 54^.

## METHODS

### Ethics patient cohort

In accordance with the Declaration of Helsinki and with the approval of the Ethics Committee of Ludwig-Maximilian-University Munich (No.: 121-09 and 20-0809) and Ethics Committe of the Technical University of Munich (No.: 635/21 S-NP) informed consent of the patients or their guardians was obtained. There was no participant compensation. Samples from a total of 31 patients with acute stroke were included into the study for multi-omic analyses. After thrombectomy, thrombi were rinsed with saline to wash away possible non-thrombus cells. Blood was collected within 90 minutes after thrombectomy in EDTA (For FACS and FACS-sort) or heparin-buffered tubes (For ScRNA-seq).

### Single-cell preparation and flow-cytometry

Thrombi immediately after harvesting were brought into single-cell suspension by gently pushing across a 70µm Cell Strainer (FALCON, Cat# 352350) into PBS (Sigma-Aldrich, Cat# D8537) and subsequently centrifuged (350rcf, 4°C, 7 minutes). For flow-cytometry based phenotyping and single-cell RNA Seq approaches, the pellet was resuspended in FACS Buffer (PBS with 0,5% BSA). The thrombus cell suspension and the patient’s blood were each mixed with the same amount of antibody mix and incubated. Erythrocyte depletion was performed by incubation with 500µl of BD Pharm Lyse Lysing Buffer (BD Biosciences, Cat# 555899), blood and thrombi were processed alike. For the flow-cytometry based monocyte and the neutrophil phenotyping, the thrombus cell-pellet as well as 200µl of the patient’s blood was each mixed with 6ml of RBC lysis buffer (containing 20g NH4Cl (1,5M), 2,5g KHCO3 (0,1M), 5ml 0,5M EDTA (10mM) and 250ml distilled water) for 10 minutes. The lysing process was stopped by addition of 6ml of cold PBS and repeated with blood after centrifugation, then samples were centrifuged and resuspended in FACS Buffer and incubated 1:1 with the respective antibody mix. For downstream 10X single-cell sequencing assays, BD Lysing buffer (Cat.: 555899) was used instead of the above-mentioned procedure for erylysis. After another centrifugation step, both samples were resuspended in FACS Buffer.

The dead cell stain was added before FACS. Flow cytometry was always performed with a LSRFortessa Flow Cytometer (BD Biosciences).

Flow-cytometry analysis was performed using FlowJo (BD). Thrombi containing mainly cell debris were not included into further analysis. For cluster identification, FlowSOM was used as described previously^35, 55, 56^, visualization was done using t-SNE.

### Preparation for prime-seq^16^ of blood and thrombus neutrophils and monocytes

Whole blood and the thrombus single-cell suspension were treated following the same erythrocyte lysis protocol as previously described for the monocyte and neutrophil FACS preparation and then stained with the respective antibody mix. After 20 minutes of incubation, the samples were washed and dead cell stain was added directly before isolating the different cell populations by FACS Sort (using the MoFlo Astrios EQ, Beckman Coulter). The isolated monocytes or neutrophils were sorted into RLT Plus Buffer with 1% β-mercaptoethanol in low-binding RNA tubes. RNA from approximately 400 sorted CD15^+^ neutrophils and 200 CD14^+^ sorted monocytes were included for the subsequent prime-RNA sequencing (prime-Seq^16^).

### prime-seq

prime-seq is a sensitive and efficient early barcoding bulk RNA-seq method described in detail in Janjic et al. (2022)^16^ and at protocols.io (dx.doi.org/10.17504/protocols.io.s9veh66). In summary, the RLT Plus Buffer (Qiagen®) with 1% βmercapto-ethanol and the included cells were further processed for library preparation. Lysate treatment was performed with Proteinase K (AM2548, Life Technologies®), isolation was performed with cleanup beads (GE65152105050250, Sigma-Aldrich®) (2:1 beads/sample ratio), and genomic DNA digestion was performed with DNase I (EN0521, Thermo Fisher®). Reverse transcription was performed with 30 units of Maxima H- enzyme (EP0753, Thermo Fisher®), and 1x Maxima H- Buffer (EP0753, Thermo Fisher®), 1 mM each dNTPs (R0186, Thermo Fisher®), 1 µM template-switching oligo (IDT), 1 µM barcoded oligo-dT primers (IDT) in a 10 µL reaction volume at 42 °C for 90 minutes. Samples from the same cell type were then pooled and cleaned using cleanup beads (1:1 beads/sample ratio). Exonuclease I (M0293L, NEB) was used to digest remaining primers following cleanup at 37 °C for 20 minutes followed by 80 °C for 10 minutes. Subsequently, the digested samples were again cleaned using cleanup beads (1:1 beads/sample ratio).

Synthesis of second strand and pre-amplification was subsequently performed using 1X KAPA HS Ready Mix (07958935001, Roche®) and 0.6 µM SINGV6 primer (IDT) in a 50 µL reaction. PCR cycles were set as follows: 98 °C for 3 minutes; 15 cycles of 98 °C for 15 s, 65 °C for 30 s, 72 °C for 4 minutes; and 72 °C for 10 minutes. Next, cleanup beads were used (0.8:1 beads/sample ratio) and then eluted in 10 µL of DNase/RNase-Free Distilled Water (10977- 049, ThermoFisher®). cDNA was quantified by the Quant-iT PicoGreen dsDNA Assay Kit (P7581, Thermo Fisher®), the distribution of size was qualified by the High-Sensitivity DNA Kit (5067-4627, Agilent®). Pre-sequencing QC was performed by Bioanalyzer traces. A total of 400 and 200 cells were included for neutrophils and for monocytes. Viability was checked during FACS-sorting, as dead cells were excluded by respective gating. The only phred based filtering in prime-seq was performed in the UMI and BC, where we had a cut off for 4 BC bases and 5 UMI bases, with a phred lower than 20. We did this to remove any data from the start that might have low sequencing quality and therefore could cause issues with demultiplexing based on the barcodes. Since the barcodes are 12 bases long, a cutoff of 4 is sufficient as anything lower would still allow a proper demultiplexing. We did not perform any filtering based on phred score of the cDNA read as part of pre-processing. Mapping Quality controls were done using the default parameters in STAR.

After quality control, cDNA was used to make libraries with the NEBNext Ultra II FS Library Preparation Kit (E6177S, NEB), using a five-fold lower reaction volume than instructed by the manufacturer. The Enzyme Mix supplied and Reaction buffer was used for fragmentation in a 6 µL reaction. Adapter ligation was performed by the supplied Ligation Enhancer, Ligation Master Mix, and an additional custom prime-seq Adapter (1.5 µM, IDT) in a reaction volume of 12.7 µL. Following ligation, double-size selection using SPRI-select Beads (B23317, Beckman Coulter®), with 0.5 and 0.7 ratios, was performed. Subsequently, sample amplification was performed using a library PCR using Q5 Master Mix (M0544L, NEB), 1 µL i7 Index primer (Sigma-Aldrich®), and 1 µL i5 Index primer (IDT) using the following setup: 98 °C for 30 s; 13 cycles of 98 °C for 10 s, 65 °C for 1 m 15 s, 65 °C for 5 m; and 65 °C for 4 m. Final double size selection was performed using SPRI-select Beads.

Concentration and quality were analysed using a high-sensitivity DNA chip (Agilent Bioanalyzer), subsequently the libraries were 150 bp paired-end sequenced on a NextSeq (Ilumina®).

### Single-cell RNA sequencing of human and mouse blood and thrombi

The Chromium Next GEM Single Cell 3‘ reagent kit v3.1 (CG000206 Rev D) from 10X Genomics® protocol was used for library preparation. To decrease batch-effect related artifacts, sample multiplexing with TotalSeqB™ anti-human or anti-mouse Hashtag Antibodies, was performed. According to the manufacturer’s instructions, first GEMs were generated, reverse transcription was performed subsequently, and cDNA was cleaned up, amplified, and size selected subsequently. After quantification and QC, gene expression libraries and cell surface libraries were constructed. The libraries were then sequenced with an Illumina NovaSeq.

### CITE-Seq

The method of Cellular Indexing of Transcriptomes and Epitopes by sequencing (CITE-Seq) has allowed us to combine the analysis of surface markers and gene expression levels for cell phenotyping. We used the following Antibodies from BioLegend: CD10 (HI10a, Cat# 312235), CD14 (clone 63D3, Cat# 367145), CD15 (clone W6D3, Cat# 323051) and CD16 (clone 3G8, Cat# 302063). Anti-CD16 antibody was employed to n=3-4 blood and thrombi, Anti CD10, CD14 and CD15 was employed to n=3 blood and thrombi, respectively.

### Bioinformatic analysis of prime-seq data

Fastqc was used to initially check the data (version 0.11.8^34^). Cutadapt (version 1.12^35^) was used to remove any regions on the 3’ end of the read where the sequence read into the poly-A tail. To filter the data, the zUMIs pipeline (version 2.9.4d, Parekh et al., 2018) was used, using a phred threshold of 20 for 4 bases for both the UMI and BC, to map the reads to the human genome with the Gencode annotation (v35) using STAR (version 2.7.3a), and count the reads using RSubread (version 1.32.4)^36, 37^. 2.7.3a), and count the reads using RSubread (version 1.32.4) ^36, 37^.

### Differential Gene Expression Analysis

Differential gene expression analysis on the bulk RNA-Seq data was performed using the poreSTAT differential expression analysis pipeline (https://github.com/mjoppich/poreSTAT/, work in progress). Differential gene expression results used in this manuscript were obtained from calling DESeq2 (version 1.30.0) with default parameters on the read counts from the zUMIs pipeline (see above). On the neutrophil bulk RNA-seq dataset 2 thrombi and 1 blood samples were removed due to too low read counts. Samples failed: 3 out of 34. Low read counts were defined as less than 140.000 reads.

### Overrepresentation analysis

Overrepresentation analysis on the prime-seq^16^ data was performed using the DAVID web interface^57, 58^.

### scRNAseq data analysis

Data were obtained and processed using CellRanger (v6.0.1 and v6.1.1) using the mm10-2020-A (for mouse data) and the GRCh38-2020-A (for human data) reference genomes. CellRanger was run using default parameters and with custom feature references. The 21053 runs were run with both antibody capture references Htag9 and Htag10 for differentiating blood and thrombus samples. The mouse data was run with antibody capture references for mouse antibodies 1-10. Finally, the remaining human samples were run with antibody capture references Htag1-Htag10 and CD10, CD14, CD15 and CD16.

The subsequent analysis was performed in R 4.0.1 with Seurat^59^ v4.0.4 (mouse) and v4.1.1 (human). For the mouse dataset, the raw Seurat counts were used and filtered such that cells have between 100 and 6000 features and at least 200 UMIs. Cells were removed if more than 15% of the UMIs per cell were of mitochondrial origin. The dataset was split according to the hashtag identified by following the HTO (Hash Tag Oligo) demultiplexing vignette of Seurat. For each library, HTO demultiplexing was performed according to the Seurat vignette with positive.quantile=0.99. Results were additionally checked using ridge plots on the HTO expression. Only cells with global identification as singlets were kept (negative and doublet cells were removed). Each sample was normalized and variable features were determined. Each sample was normalized using Seurat’s NormalizeData function and variable features were determined – as the standard analysis pipeline from the SatjaLab. Human cell-cycle genes from Seurat were converted to mouse using the biomart^60^ library. The samples were scaled, regressing our percent.rp, percent.mt, nCount_RNA as well as S.Score and G2m.Score from Seurat’s CellCycleScoring function. A total of 61.943 cells were included for subsequent analyses. The samples were then integrated using rpca with 3000 anchor features and 50 PCs. The integrated object was scaled, and PCA was run. UMAP and neighbours were identified from the first 50 PCs. Clusters were identified at a resolution of 0.2 . Differential gene expression was calculated using the t test implemented in Seurat.

The human datasets were loaded into Seurat objects in a library-wise pattern from the raw cellRanger output. The samples were filtered such that cells have at least 100 UMIs and between 100 and 6000 features in order to capture also neutrophils, which have little RNA expression. For each library, HTO demultiplexing was performed according to the Seurat vignette. After HTO demultiplexing, the libraries were each normalized and variable features were detected. A total of 4000 integration features was selected for scaling the libraries regressing out percent.rp, percent.mt, nCount_RNA as well as S.Score and G2m.Score from Seurat’s CellCycleScoring function, which was also run on the libraries. Integration itself was performed for all libraries using 3000 integration features and following the reciprocal PCA integration vignette using 30 PCs and 10 anchors for integration. After integration, the antibody capture for the CITE-seq counts was prepared. For each library, the counts were subset such that the normalization using the CLR method with margin 2 was performed on only those antibodies that were measured in the respective samples. According to the recommendations of the Seurat authors, CLR normalization was conducted with margin 2, because both the hashtag oligonucleotides as well as the CITE-seq antibodies are competing in binding to the GEM, and thus CITE-seq antibodies could not be guaranteed to attach to saturation (which would be the assumption using margin 1 normalization). In order to add the assay to the integrated object, the count matrix was extended with zeros for all remaining cells for all antibodies. There is no further integration of the CITE-seq counts required, because CITE-seq counts were not used for integrating the samples, nor for dimensionality reduction. In analogy to the RNA assay, any expression analysis is performed on the unscaled per-cell-normalized ADT values. Heatmaps were created using the ComplexHeatmap package on the scale.data slot of the RNA assay of the scRNA-seq data^61^.

We performed condition-wise gene module detection using the method and script provided by Kazer et al^26^. In brief, this method takes the gene expression values of the genes represented by the first few principal components of the scRNA-seq object as input for WGCNA functions. Here, we chose the maximal 500 genes reported by the PCASigGenes function of Seurat for each of the first eight PCs. The resulting adjacency matrix is transformed into a TOM and hierarchically clustered. These clusters are merged if not too dissimilar. After testing the modules for their significance (p-Value threshold of 0.05, 10 bins and 100 permutations), the remaining modules are tested regarding the temporal variation (sample size is minimum of module size and 150, 1000 tests, order of conditions Blood < Thrombus, p-value threshold of 0.05). The remaining modules are added to the Seurat object using the AddModuleScore function (ctrl set to 5) and are reported for further visualization and discussion. All samples were processed using 100 genes from each of the 10 principle components. All samples were processed with a soft power of 7, the monocytes with a soft power of 10.

scRNA-seq data will be accessible via Zenodo (https://doi.org/10.5281/zenodo.10466854).

### Chemokine interactome analysis

Chemokine ligand-chemokine receptor interactions were collected from two different resources^62^ and https://www.rndsystems.com/pathways/chemokine-superfamily-pathway-human-mouse-lig-recept-interactions (year of access: 2021). Interactions are either classified as antagonist, agonist or undefined. In general, the steps described by Armingol et al.^63^ are followed for determining cell-cell communications. The experimental expression data (for each cluster) is read in and filtered to only contain genes from the above collection of chemokine interactors. For each ligand-receptor pair, and for each cluster-pair, the communication score is calculated. This communication score is the product of the ligand expression and the receptor expression (expression product).

This results in a data frame in which for each ligand-receptor pair in each cluster pair a score is associated. To determine the total communication between two clusters, all communication scores between these clusters are summed.

In a second step, the data frame is arranged into matrix form, keeping only clusters of interest (or all, if no filtering was requested). The chord diagram (taken from https://github.com/tfardet/mpl_chord_diagram) displays the accumulated LR-interactions between clusters (all, or only for monocytes and neutrophils).

In addition, the matrix plot shows the scaled (z-score) expression scores for all interactions in the selected clusters. In a filtered version, only interaction which have at least in one cluster pair a z-score > 1 are shown. Finally, the chemokines overview displays the ligand-receptor map in the lower left corner, and shows the expression values for the receptors in the selected clusters on top, and those for the ligands to the right. This visualization allows for a concise overview of ligand and receptor expressions, together with an overview of possible LR- interactions. Source code for this analysis is available online https://github.com/mjoppich/chemokine_interactionmap.

### scVelo

For the velocity analysis, the barcodes of all cells in the integrated Seurat object were written to disk per library. This list of barcodes is important for running velocyto^64^ if also barcodes are used that are not part of the filtered barcodes from cellRanger. Apart from specifying the barcodes of the required cells, velocyto was run with default parameters. After performing the couting of unspliced RNAs using velocyto, the acquired loom files were matched with the integrated Seurat object and the unspliced count matrix is written to disk together with a loomified version of the integrated Seurat object and additional meta information (e.g. UMAP coordinates). In the second step, this object is read using scvelo^31^.

The UMAP embedding is taken over and spliced (cellRanger counts) and unspliced (velocyto) counts are stored in their respective layers. Genes that were not present in at least 50 cells were filtered out and the remaining matrix was normalized. 3000 highly variable genes were called using the Seurat-flavor before calculating PCs. In order to have a functioning neighbour graph, bbknn was called with batch_key identifying the single libraries (matching the integration strategy from Seurat). The general scvelo vignette for performing velocity analyses was followed to calculate velocity in deterministic mode.

The source code of the scRNA-seq data analysis will be made available together with the preprocessed Seurat objects upon publication.

### Neutrophil and PBMC isolation and subsequent pharmacological hypoxia mimicking

For further examination of neutrophil granulocytes we isolated granulocytes with the EasySep® Direct Human Neutrophil Isolation Kit (Stemcell Technologies, Cat# 19666), following the instructions provided by the manufacturer. Isolated neutrophils were normalized to a concentration of 1 million cells/ml. PBMCs were isolated from whole blood with BD Vacutainer CPT tubes (BD Biosciences, Cat# 362780) and adjusted to a cell concentration of 10 million cells/ml.

The respective cells were distributed into 14ml Polystyrene Round-Bottom Tubes (Falcon, Cat# 352057). After centrifuging the tubes (350rcf, 4°C, 5 minutes) half of them was resuspended in a medium containing RPMI, 2mM L-Glutamin and 2% FBS, the other half in the same medium with additionally 50µM Roxadustat (Selleckchem, Cat# S1007) and incubated for 16 to 18 hours (37°C, 5% CO2). Then, after another centrifugation and resuspension step, the samples were stained with the antibody mix and washed again.

Directly before FACS or FACS Sort, dead cell stain was added.

Non-classical and classical monocytes from the incubated PBMCs were FACS-sorted with a BD FACSMelody™ Cell Sorter into RLT Plus Buffer with 1% β-mercaptoethanol and subsequently analysed by prime-seq^16^, neutrophils were analysed with a LSRFortessa Flow Cytometer (BD Biosciences).

### Artificial coagulation activation and subsequent monocyte analysis

We used citrate coated tubes to take blood from healthy donors, which was afterwards filled into two Polystyrene Round-Bottom Tubes. After adding thrombin and calcium-chloride to one of the tubes and incubation for 4 hours, the coagulum was pushed through a 70 µm mesh, solved in PBS, centrifuged (350rcf, 7 minutes) and resuspended in FACS Buffer. Afterwards, the cell suspension from the coagulum and the blood from the other tube were incubated with the antibody master-mix. For the erythrocyte lysis, BD Pharm Lyse (BD Biosciences, Cat# 555899) was used. Directly before the Sorting process, dead cell stain was added.

FACS Sort was performed with the BD FACSMelody™ Cell Sorter, classical and non-classical monocytes were FACS-Sorted.

### Thrombolysis assay

After taking blood from healthy donors, we isolated neutrophils and plasma from lithium-heparin-coated tubes and platelet-poor plasma from citrate-coated tubes. The coating of the Ibidi® µ-slides IV 0,4 Uncoated (Cat# 80601) with fibrin was performed by adding PBS mixed with recombinant human albumin (Sigma-Aldrich, Cat# A9731), fibrinogen from human plasma AF488 Conjugate (Thermo Fisher Scientific, Cat# F13191), platelet-poor plasma, 100mM calcium-chloride and thrombin from bovine plasma (Sigma, Cat# T4648-1KU) to the chambers. After 15 minutes, the slides were washed three times with PBS.

After staining the isolated neutrophils with the Cell Proliferation Dye eFluor™ 670 (Invitrogen, Cat# 65-0840-85), 2x10^5^ neutrophils were added to the chambers, followed by a short centrifugation step (40rcf, 7 minutes). Finally, after washing the chambers three times with PBS, a medium consisting of RPMI with 2mM L-Glutamin, 20% human plasma and for the treated chambers 50µM Roxadustat was added and the chambers were incubated at 37°C and 5% CO_2_ for 16 hours.

Per donor, three chambers with Roxadustat treated neutrophils and three chambers with untreated neutrophils were analysed. Per chamber, five pictures with 20x augmentation were taken with the Zeiss Axiocam 506 mono microscope and subsequently analysed with the Fiji ImageJ software.

### Deep vein thrombosis - Mouse model of flow restriction in the IVC^4^

After a median laparotomy the IVC was exposed and a space holder was positioned followed by a narrowing ligature. Subsequently, the wire was removed to avoid complete vessel occlusion. Side branches were not ligated or manipulated. All groups were age, sex, and weight matched. Mice with bleedings or any injury of the IVC during surgery were excluded from further analysis. For thrombus weight measurement after 48 hours, the IVC was excised just below the renal veins and proximal to the confluence of the common iliac veins.

### Intravital neutrophil fate mapping

Directly before the flow restriction of the IVC, FITC anti-mouse Ly-6G antibody was injected i.v. into the mouse tail vein (5µg/25g mouse, Clone 1A8, Biolegend). 48 hours after the first FITC anti-mouse Ly-6G antibody injection, PE anti-mouse Ly-6G Antibody was injected i.v. into the tail vein of the same mice (5µg/25g mouse, Clone 1A8, Biolegend). 24 hours after the second injection (72 hours after the restriction of the IVC), the IVC was collected, embedded in OCT and stored at -80°C for histology.

### Bone marrow transplant

Bone marrow chimeras were generated by irradiation of recipient C57Bl6 mice. Six- to eight-week-old mice were irradiated twice within a 4-h interval (195 kV, 10 mA) and received isolated bone marrow from control C57/Bl6 mice or Nr4a1^-/-^ mice on C57/Bl6 background (The Jackson Laboratory, Cat.: 006187).

### Immunofluorescence staining and quantification of thrombus neutrophils in mouse thrombi*

The IVC was embedded in OCT, stored at -80°C and cut with a cryotome (CryoStar NX70 Kryostat, ThermoFisher Scientific). Samples were fixed with 4% formaldehyde and blocked with the respective serum. In Nr4a1^-/-^ and Nr4a1^+/+^ mouse thrombus sections neutrophils were visualized by anti-Ly6G (clone: 1A8; isotype: rat IgG2a) and MPO (rabbit polyclonal, #GA51161-2, DAKO; isotype: rabbit immunoglobulin fraction, DAKO). Alexa labeled secondary antibodies (Invitrogen) were used for detection. DNA staining was performed using Hoechst (Invitrogen, Cat. # H3570). Finally mounting medium (DAKO) was applied to place a cover slip. The image acquisition was carried out by using a Zeiss Axio imager microscope with an AxioCam. After image acquisition, cells were manually counted. For quantification of NETS three criteria had to be fulfilled: (1) Presence of extracellular DNA protrusions, (2) the protrusion has to originate from a Ly6G-marked cell, (3) the DNA-structure has to be decorated by MPO. To visualize neutrophils and urokinase expression in human thrombus, we used Hoechst dye (Invitrogen, Cat. # H3570), anti-MPO (R&D systems, Cat. # AF3667), and anti-Urokinase (Abcam, Cat. # ab24121). The imaging was performed by the ZEISS LSM880 (Confocal, Serial # 3851003082) Axio Imager.M2 (Carl Zeiss Microscopy GmbH, Göttingen).

### ROTEM Analysis

We used rotational thromboelastometry (ROTEG-05, Pentapharm GmbH, Munich, Germany) to measure the different coagulative capacities of murine whole blood from wildtype and Nur77-/- mice under low sheer stress conditions, comparable to those in the lower Vena cava [103]. Murine blood was drawn into citrate tubes (#04.1952.100, SARSTEDT AG & Co. KG, Nuembrecht, Germany) immediately analyzed. Therefore, we mixed 300 µl of murine whole citrate blood with 20 µl StarTEG solution (#503-10, Tem Innovations GmbH, Munich, Germany) and 20 µl EXTEM solution (#503-05, Tem Innovations GmbH, Munich, Germany) in reagent cups (#200011, Tem Innovations GmbH, Munich, Germany) and measured clotting time (CT), clot formation time (CFT), maximum clot firmness (MCF) and maximum lysis (ML) at 37 °C ^65^.

### Antibody panels*

*In the Roxadustat *in vitro* experiment group, CD87 in PE (Biolegend, Cat.: 336906) and CD177 in PE-Dazzle (Biolegend, Cat.: 313226) were used for analysis.

| Color | Antigen | Company | Cat. # |
| --- | --- | --- | --- |
| BV650 | CD45 | BioLegend | 304044 |
| FITC | CD14 | BD<br>Pharmingen | 557153 |
| PERCP-Cy5.5 | Siglec 8 | BioLegend | 347107 |
| APC | CD15 | BioLegend | 301908 |
| APC-Cy7 | CD19 | BioLegend | 363010 |
| BV605 | HLA-DR | BioLegend | 307640 |
| BV785 | CD123 | BioLegend | 306032 |
| BV510 | CD3 | BioLegend | 317332 |
| BV711 | CD56 | BioLegend | 362542 |
| AF700 | CD34 | BioLegend | 343622 |
| Pacific Blue | CD41 | BioLegend | 303714 |
| SytoxOrange | Nucleic acid<br>stain | Invitrogen | S11368 |

FACS Neutrophil Panel\*:
| Color | Antigen | Company | Cat. # |
| --- | --- | --- | --- |
| BV650 | CD63 | BioLegend | 353026 |
| FITC | CD177 | BioLegend | 315804 |
| PE | VEGFR-1 | MACS Miltenyi<br>Biotec | 130-124-438 |
| PERCP-Cy5.5 | TCR $\alpha/\beta$ | BioLegend | 306724 |
| APC | CD15 | BioLegend | 301908 |
| APC-Cy7 | CD184 | BioLegend | 306528 |
| BV605 | CD49d | BioLegend | 304324 |
| BV785 | CD62L | BioLegend | 304830 |
| BV510 | CD16 | BioLegend | 302048 |
| BV711 | CD11b | BioLegend | 301344 |
| AF700 | IL-17A | BioLegend | 512318 |
| PE-Dazzle | CD31 | BioLegend | 303130 |
| PE-Cy7 | CD284 | Invitrogen | 25-9917-42 |
| SytoxBlue | Dead cell stain | Invitrogen | S34857 |

\*In the Roxadustat *in vitro* experiment group, CD87 in PE (Biolegend, Cat.: 336906) and CD177 in PE-Dazzle (Biolegend, Cat.: 313226) were used for analysis.

FACS Monocyte Panel:
| Color | Antigen | Company | Cat. # |
| --- | --- | --- | --- |
| BV650 | CD45 | BioLegend | 304044 |
| FITC | CD14 | BD<br>Pharmingen | 557153 |
| PE | CD93 | BioLegend | 336108 |
| PERCP-Cy5.5 | CD36 | BioLegend | 336224 |
| APC | SLAN | MACS Miltenyi<br>Biotec | 130-117-919 |
| APC/Fire 750 | IgE | BioLegend | 325516 |
| BV605 | HLA-DR | BioLegend | 307640 |
| BV786 | CD192 | BioLegend | 357234 |
| BV510 | CD16 | BioLegend | 302048 |
| BV711 | CD86 | BioLegend | 305440 |
| AF700 | CD11c | BioLegend | 337220 |
| PE-Dazzle | CD11a | BioLegend | 301232 |
| PE-Cy7 | CD9 | BioLegend | 312116 |
| SytoxBlue | Dead cell stain | Invitrogen | S34857 |

FACS Sort Panel for Neutrophils and Monocytes:
| Color | Antigen | Company | Cat. # |
| --- | --- | --- | --- |
| FITC | CD45 | BioLegend | 368508 |
| PB | CD14 | BD Biosciences | 558121 |
| PE-Cy7 | CD16 | BD Biosciences | 557744 |
| PERCP-Cy5.5 | Siglec 8 | BioLegend | 347107 |
| APC | CD15 | BioLegend | 301908 |
| APC-Cy7 | CD19 | BioLegend | 363010 |
| BV510 | CD3 | BioLegend | 317332 |
| AF700 | CD34 | BioLegend | 343622 |
| SytoxOrange | Dead cell stain | Invitrogen | S11368 |

FACS Sort Panel for Non classical and Classical Monocytes:
| Color | Antigen | Company | Cat. # |
| --- | --- | --- | --- |
| FITC | CD14 | BD Pharmingen | 557153 |
| PE-Cy7 | CD16 | BD<br>Biosciences | 557744 |
| APC-Cy7 | CD56 | BioLegend | 362512 |
| PE | CD3 | BioLegend | 300308 |
| PE | CD20 | BioLegend | 302306 |
| SytoxRed | Dead cell<br>stain | Invitrogen | S34859 |

\*Not every single antibody used for staining was shown or used for analysis.

## AUTHORSHIP CONTRIBUTION

Conception, experimental design, project administration and supervision: K.P., L.N. K.S.; Writing original draft: K.P.; Patient identification, patient information and informed consent acquisition, sample acquisition: N.M., M.R.H.P., T.B.B., T.L., S.T.; Methodology, investigation and formal analysis: K.P., L.N., M.J., S.B., V.K., L.E., A.M.N., R.K., N.M., B.K., S.S., A.J., V.P., A.D.S., M.V.; Writing & editing, all authors; Visualization: K.P., A.L., M.J.; Funding acquisition: K.P., S.T., M.D., S.T., B.E., R.Z., S.M., L.N., K.S.

## Suppl. Figure captions

**Suppl. Fig. 1.**
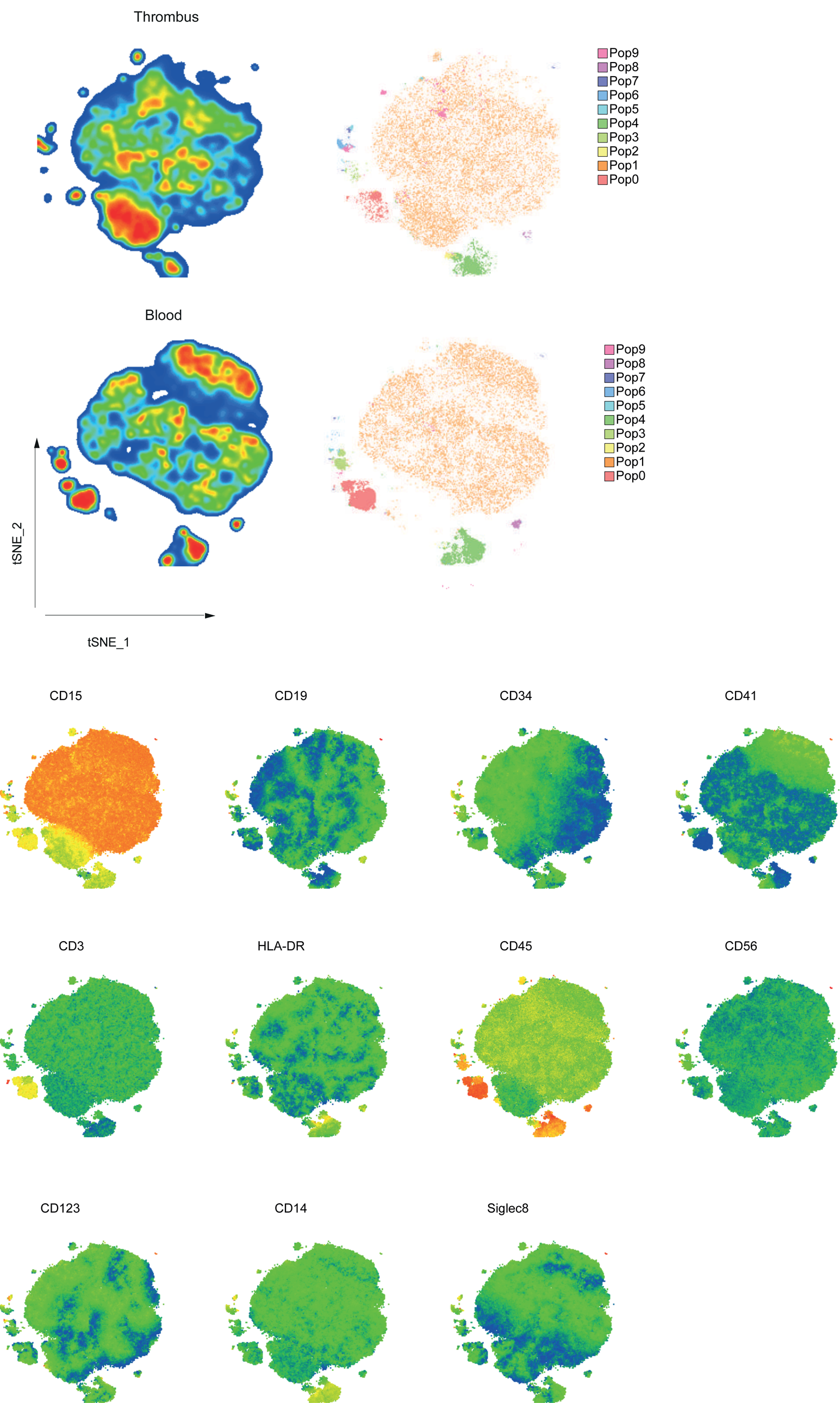
Multicolour flow cytometry and subsequent dimensionality reduction with tSNE and FlowSOM based cluster identification in blood and thrombus as one concatenated file (top). tSNEs with expression for individual surface makers of blood and thrombus cells included (bottom).

**Suppl. Fig. 2.**
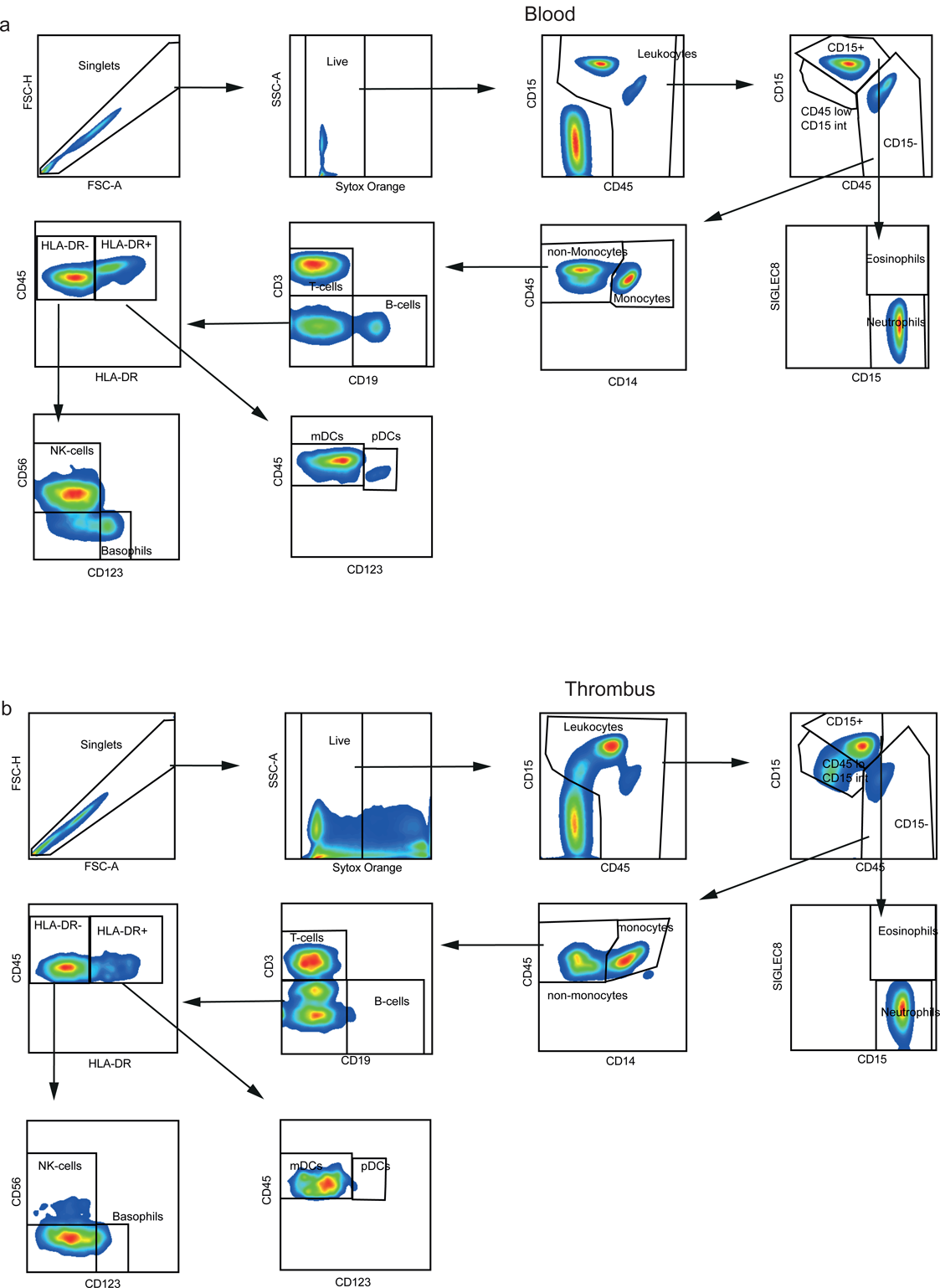
Gating strategy, allowing flow-cytometric quantification of all major immune-cells withing thrombus and blood.

**Suppl. Fig. 3.**
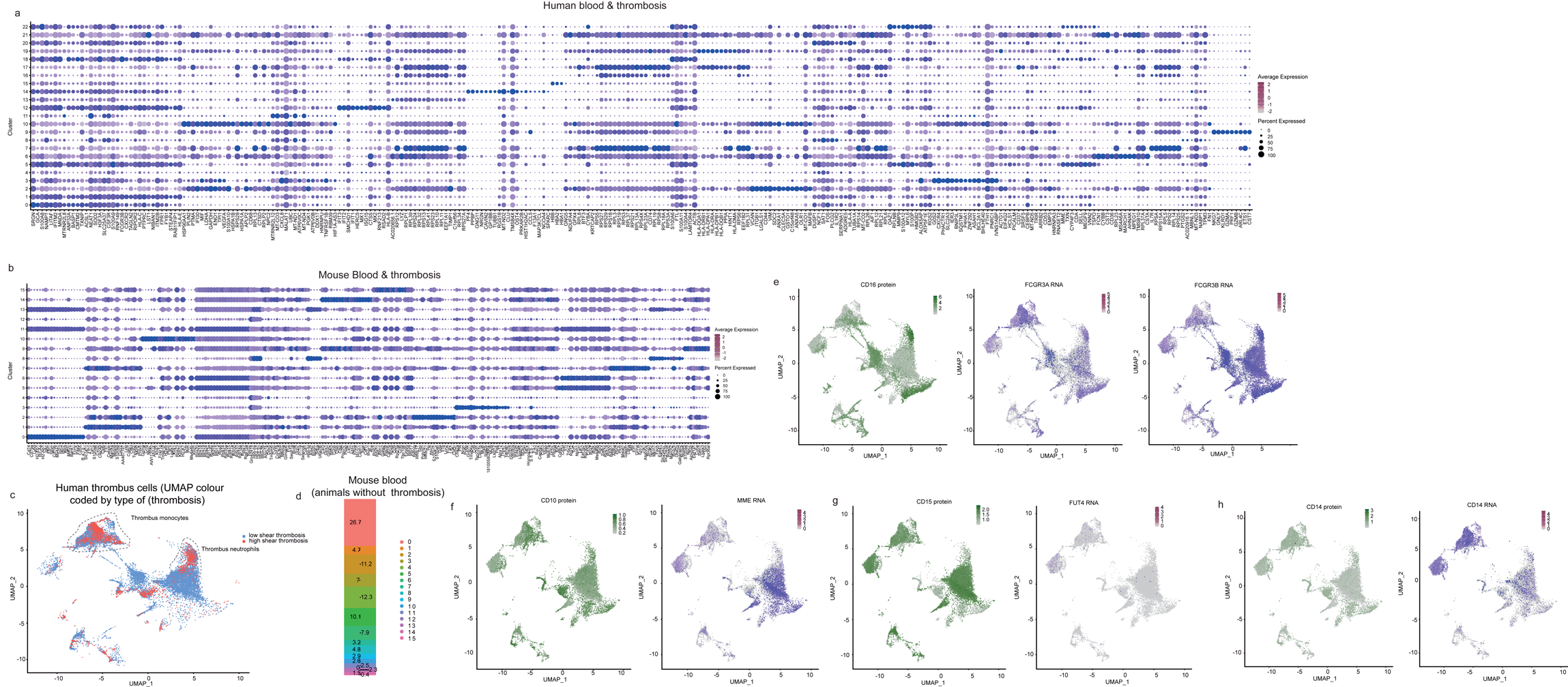
**(a)** Dot plot depicting marker genes of all clusters in human thrombosis and blood further described in Figure 1 selected by maximizing pct.1 * logFC. **(b)** Dot plot depicting marker genes of all clusters in murine thrombosis and blood further described in Figure 1, selected by maximizing pct.1 * logFC. **(c)** UMAP of human blood and thrombus scRNA-seq data, coloured by type of thrombosis (blue depicts blood and thrombus cells from low-shear, red from high-shear thrombosis), n=2 high shear thrombi, n=4 low shear thrombi. **(d)** Plot depicting blood leukocytes from mice without thrombosis (in contrast to thrombus and matched blood from same mice in Fig. 1l Barplot). Cells from n=3 mice. (**e**)-(**g**) CITE-Seq feature plots depicting gene and protein expression of CD16 (**e**), CD10 (**f**), CD15 (**g**) and CD14 (**h**) in n=3- 4 human thrombi and blood, respectively.

**Suppl. Fig. 4.**
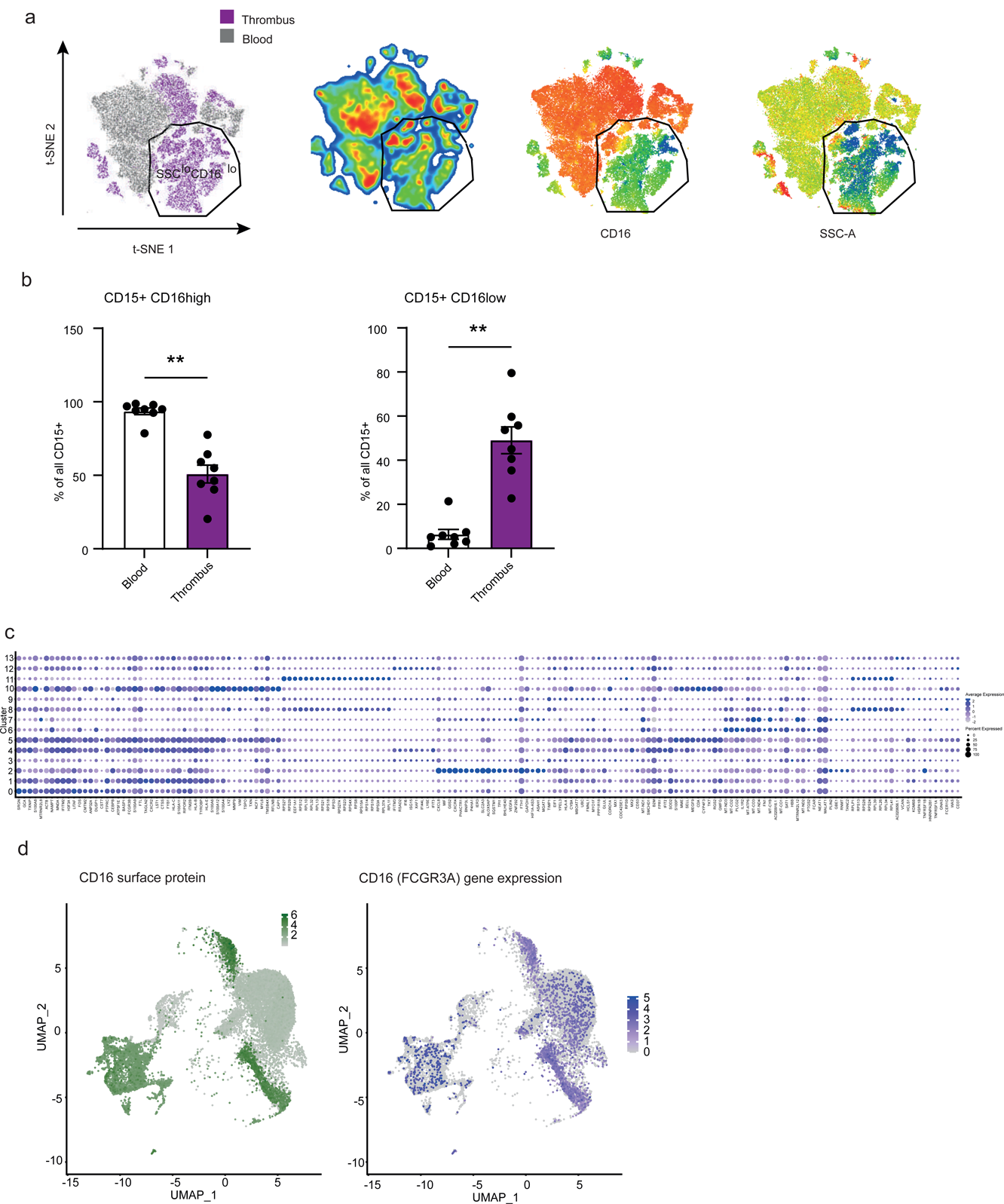
Flow-cytometry based phenotyping of thrombus and blood neutrophils. **(a)** tSNE based dimensionality reduction of blood and thrombus neutrophils with subsequent colour coded illustration by origin (left), cell frequencies relative to all neutrophils (middle), and feature plot depicting expression levels of CD16 and SSC-A levels (right) (blue depicts low expression or relative cell abundance, green intermediate and red depicts high expression or relative cell abundance, respectively). **(b)** Flow-cytometry based quantification of CD15^+^ CD16^high^ and CD15^+^ CD16^low^ neutrophils in blood and thrombi. n=8 blood and n=8 thrombi. Wilcoxon matched-pairs signed rank test was used for non-parametric distributed data. ** P<0.01. **(c)** Dot plot depicting neutrophil cluster defining genes of all clusters in human thrombosis and blood further described in Figure 2 selected by maximizing pct.1 * logFC. **(d**) Feature plot of Single-cell CITE-Seq of blood and thrombus neutrophils showing CD16 (FCGR3A) gene and CD16 protein expression. n=4 thrombi and n=3 matched blood.

**Suppl. Fig. 5.**
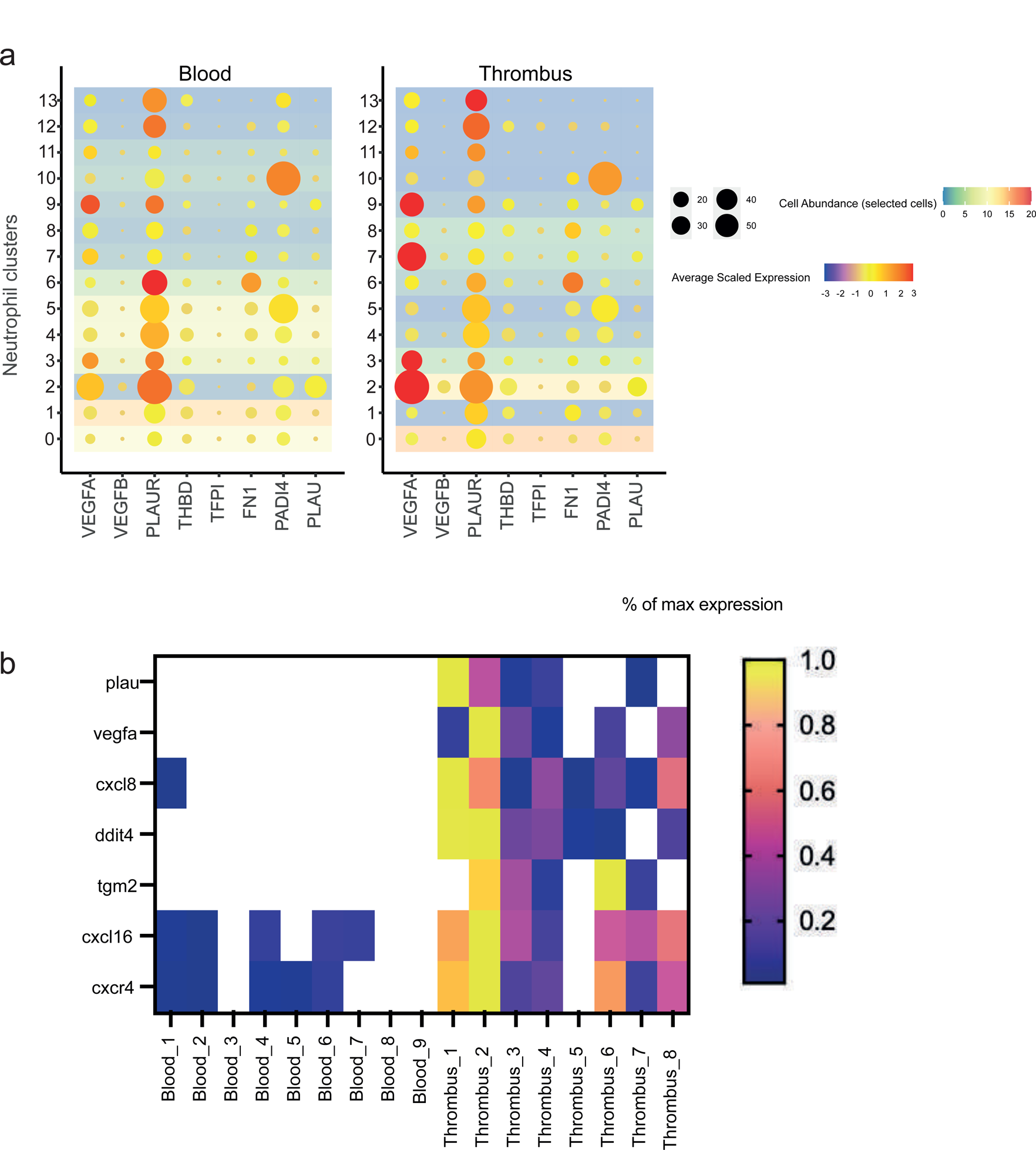
**(a)** Dotplot depicting functional neutrophil genes with direct thrombus modulating capacities. Color of the dot depicting gene expression levels, size of the dot depicts relative amount of cells within the cluster actually expressing the gene, background colour depicts relative frequency of the cell cluster within the respective tissue (blood or thrombus). **(b)** Heatmap of key differentially regulated genes between blood and thrombus neutrophils as determined by prime-seq^16^ of FACS-Sorted neutrophils. n=8 thrombi, n=9 blood (independent cohort).

**Suppl. Fig. 6.**
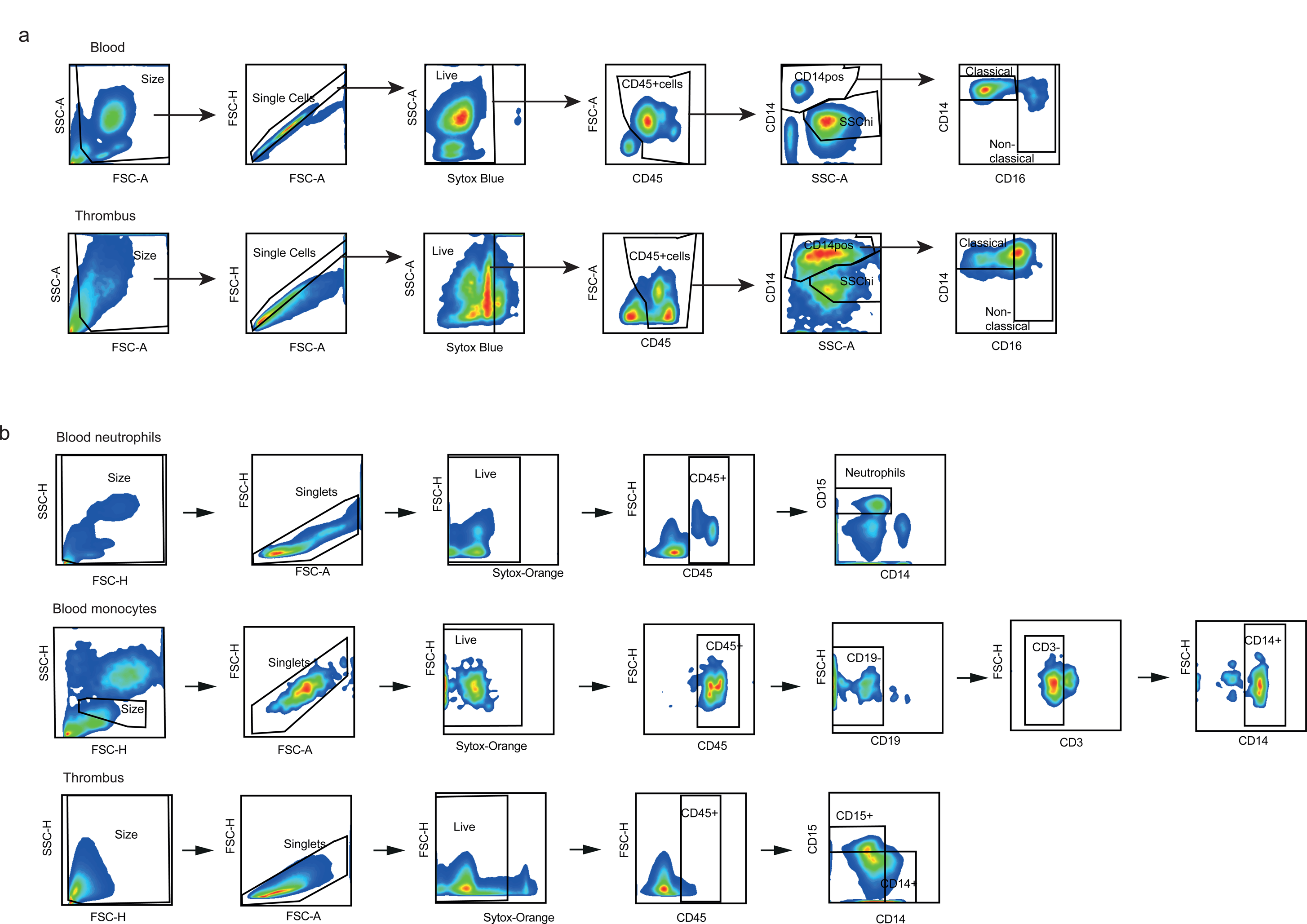
**(a)** Flow-cytometry based gating strategy for classical and non-classical monocytes. **(b)** FACS-Sort gating strategy of blood and thrombus monocytes and neutrophils.

**Suppl. Fig. 7.**
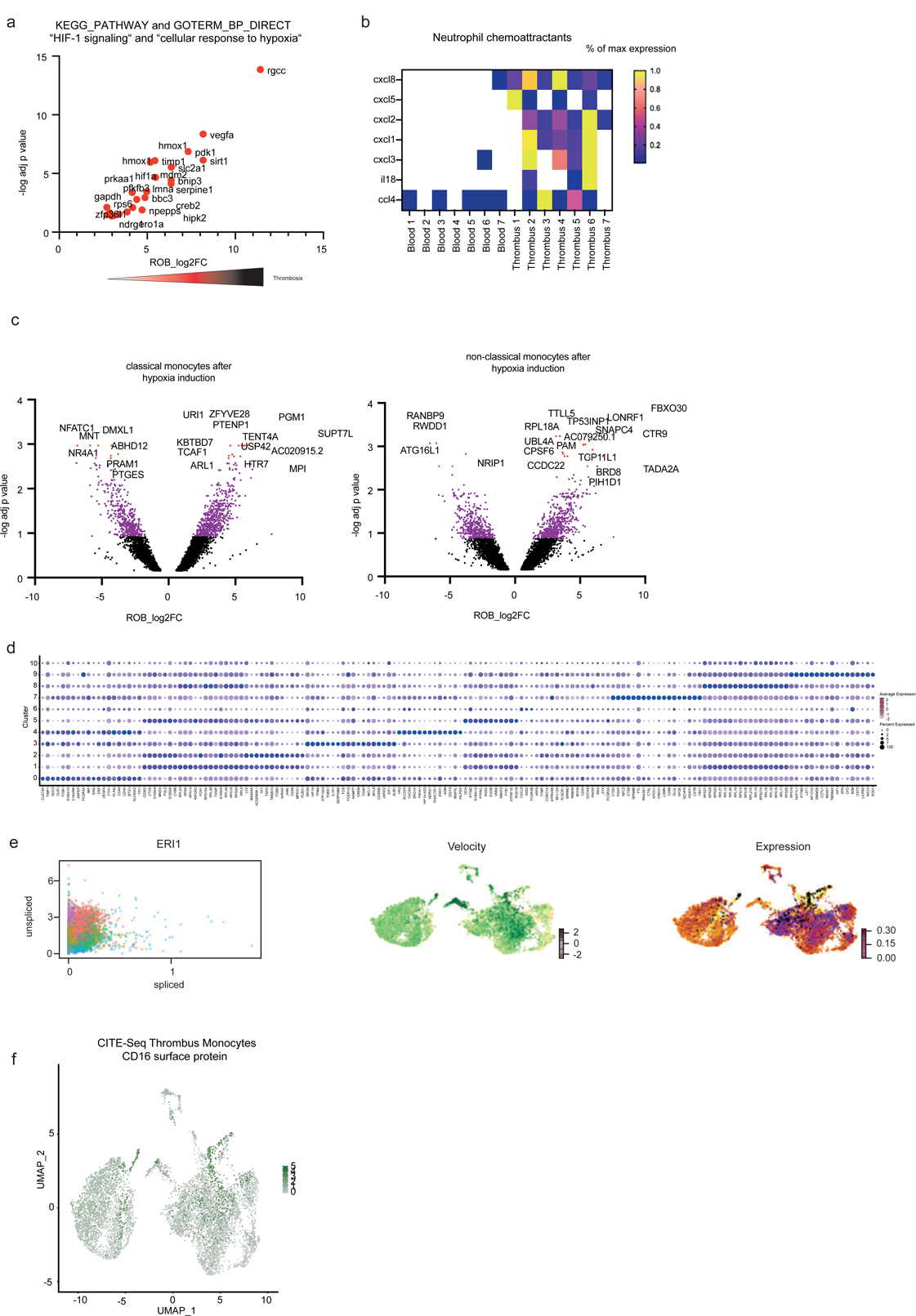
**(a)** Genes involved in enriched KEGG_Pathway and GO_BP_Direct Term: “HIF- 1 signaling“ and “cellular response to hypoxia“. **(b)** Heatmap of thrombus monocyte genes depicting expression of neutrophil chemoattractants as analysed by prime-seq^16^ of FACS/Sorted monocytes in thrombus and blood. **(c)** Volcano plots of differentially regulated genes as determined by prime-seq^16^ of FACS-Sorted classical and nonclassical monocytes after Roxadustat mediated HIF1α stabilization in monocytes. n=5/group. **(d)** Cluster defining genes of subclusters depicted in Figure 5 (sorted by pct1 * logFC). **(e)** scVelo^31^ based analysis of spliced and unspliced ERI1 expression (coloured by cluster, left) and the calculated velocity (middle) as well as expression (right) as depicted by feature plots. **(f)** Single-cell CITE-seq depicting CD16 expression on protein level. n=4 thrombi and n=3 matched blood.

**Suppl. Fig. 8.**
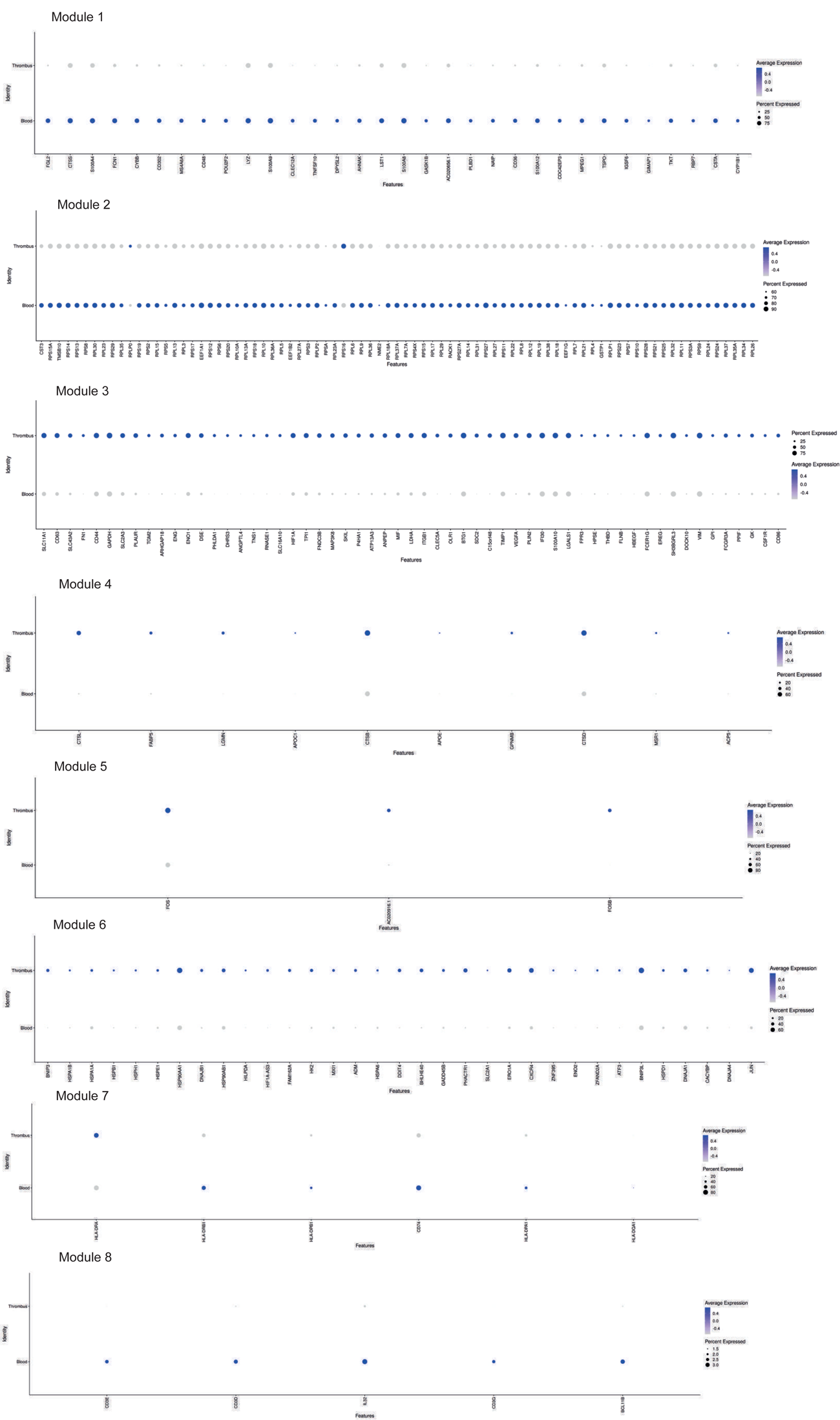
Identified Gene Modules in monocytes (Selected genes of Module 3 are also depicted in Main Figure 3) and respective genes included in the respective module with its expression as depicted by dot plots (see methods for further explanation of WGCNA analysis).

**Suppl. Fig. 9.**
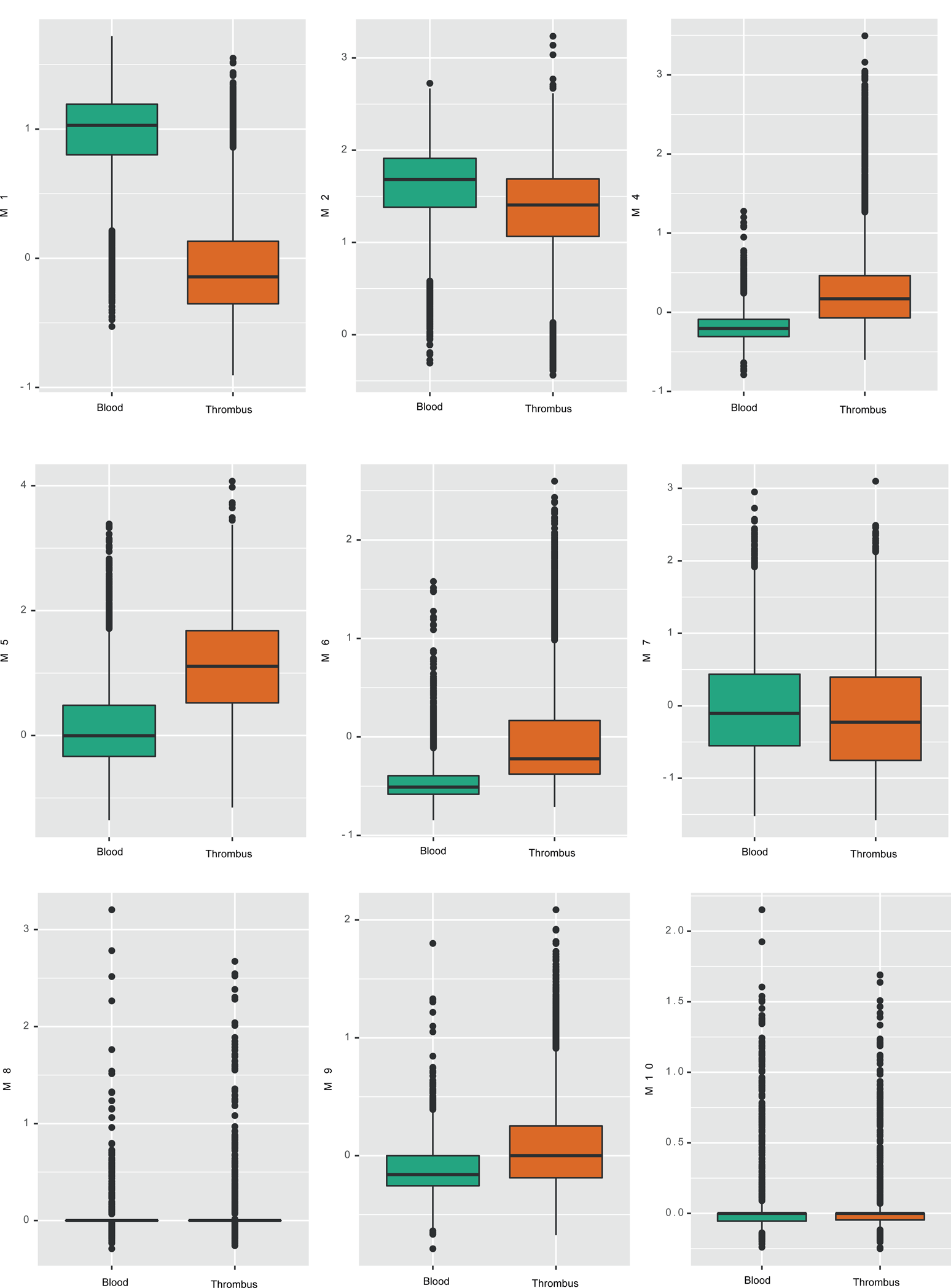
Module Score of all WGCNA Gene Modules in monocytes depicted across blood and thrombus, illustrated as box-plots showing median and interquartile ranges in blood and thrombus monocytes.

**Suppl. Fig. 10.**
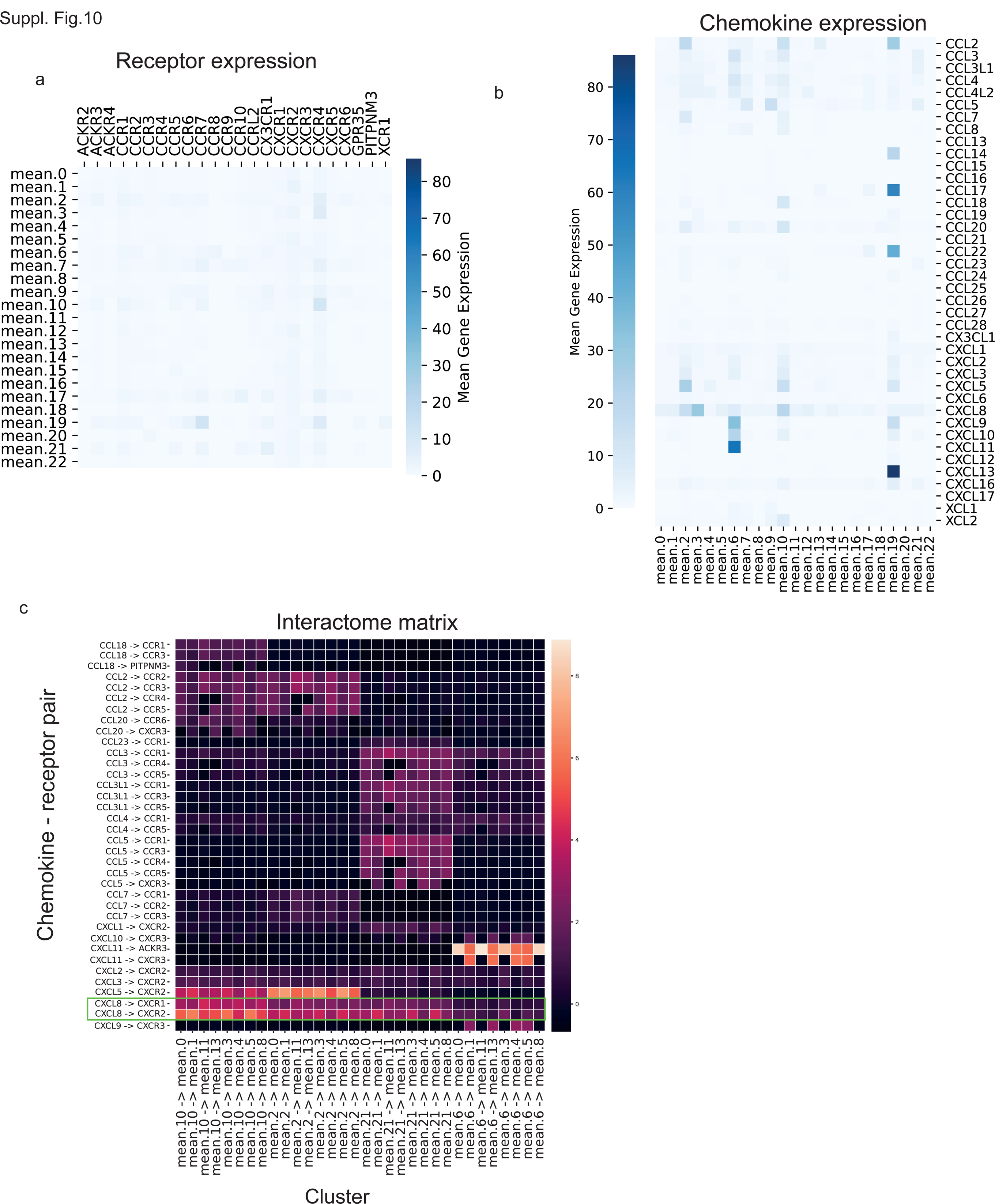
Cell clusters further described in Figure 1k, communication pathways summarized in Figure 4a. **(a)** ChemokineReceptor expression across clusters as depicted by a heatmap. **(b)** Chemokine (ligand) expression across clusters as depicted by a heatmap. **(c)** Chemokineligand-receptor interactome matrix based on **(a)** and **(b)**. Arrows pointing from source clusters (ligand expressed) to target clusters (receptor expressed). Interactome matrix focuses on blood and thrombus innate immune cells.

**Suppl. Fig. 11.**
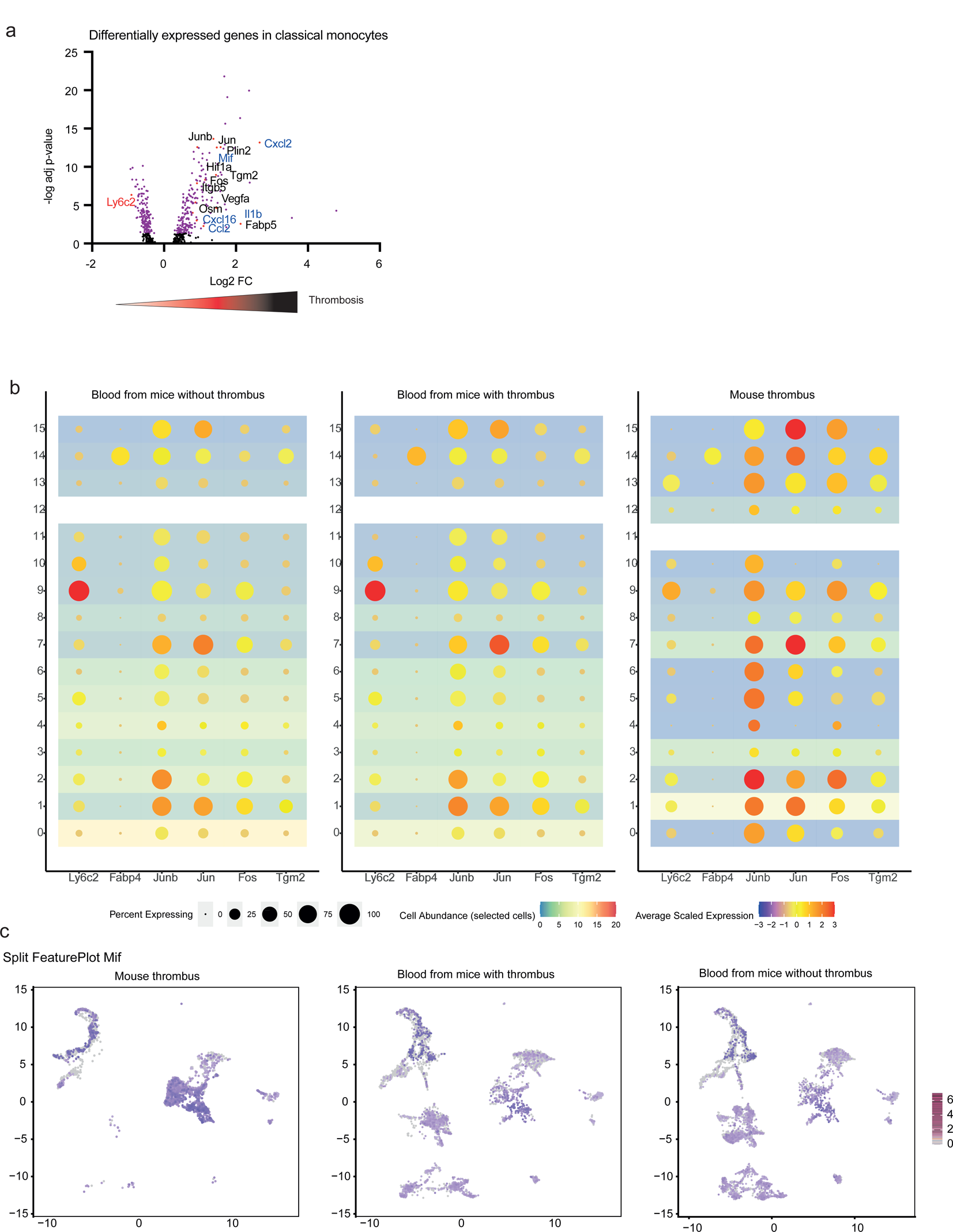
**(a)** Volcano plot depicting differentially regulated genes in mouse classical monocytes between thrombus and matched blood. Blue depicts upregulated neutrophil chemoattractants in monocytes. Red depicts downregulated Ly6c2 gene, representing a less classical phenotype. **(b)** Monocyte clusters in murine thrombi and blood, depicting changes in similar genes as previously described for human thrombus monocytes. **(c)** Representative feature plot depicting the changes between thrombus and blood monocytes in respect to the expression of the neutrophil chemoattractant MIF.

**Suppl. Fig. 12.**
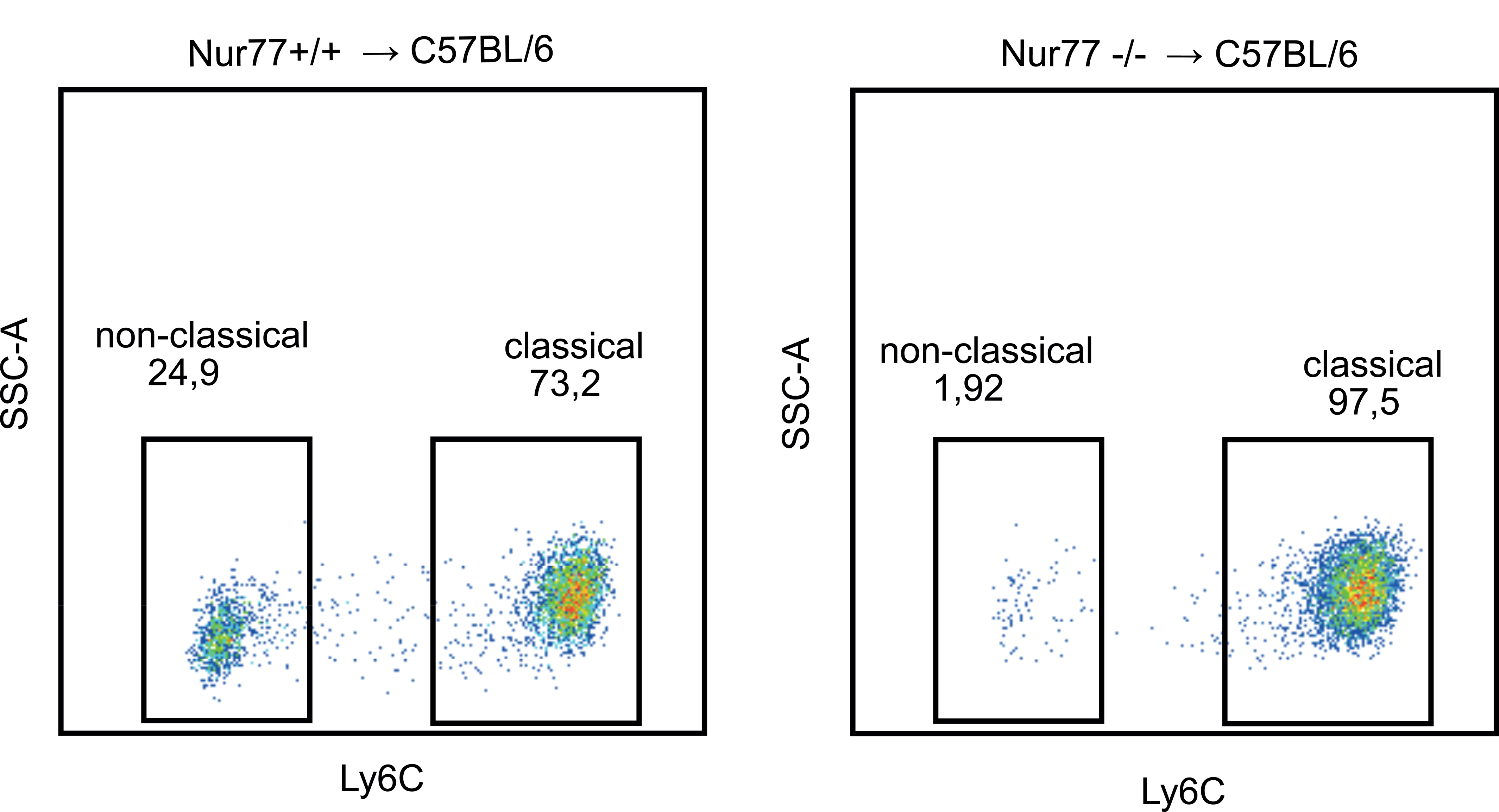
Representative gates depicting the classical and non-classical monocyte ratio in the blood of Nr4a1^+/+^ and Nr4a1^-/-^ bone marrow chimeras.

**Suppl. Table 1.**
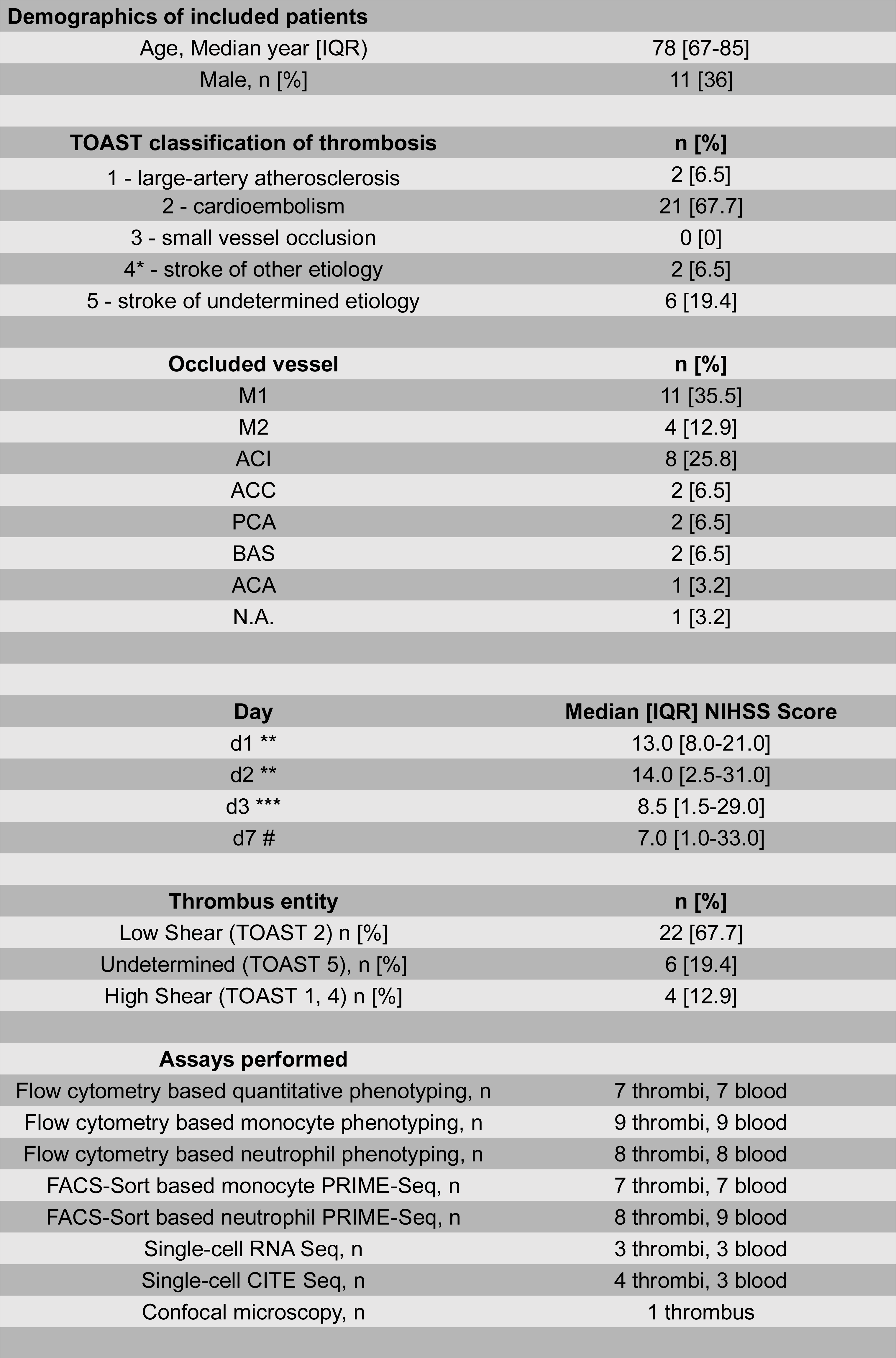
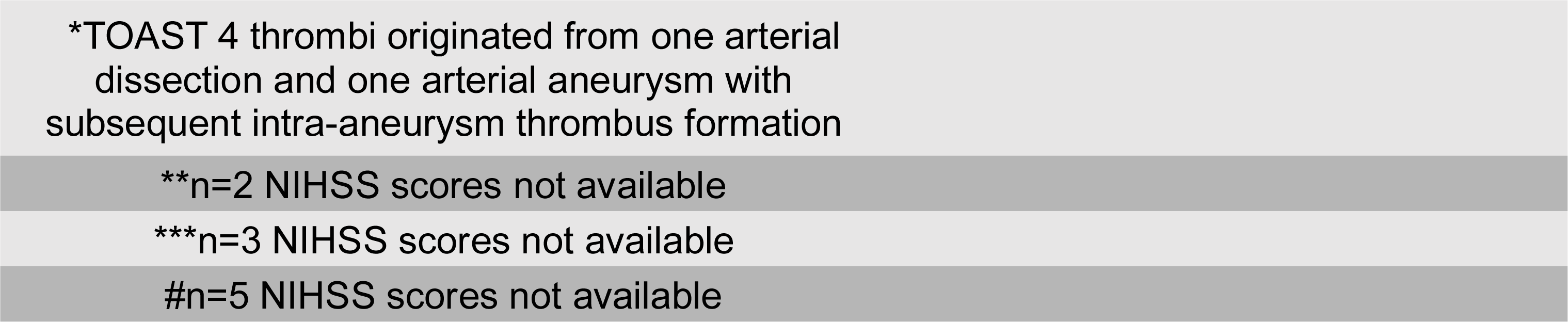
Human biosamples.

| **Demographics of included patients** |  |
| --- | --- |
| Age, Median year [IQR] | 78 [67-85] |
| Male, n [%] | 11 [36] |
| **TOAST classification of thrombosis** |  |
| 1 - large-artery atherosclerosis | 2 [6.5] |
| 2 - cardioembolism | 21 [67.7] |
| 3 - small vessel occlusion | 0 [0] |
| 4* - stroke of other etiology | 2 [6.5] |
| 5 - stroke of undetermined etiology | 6 [19.4] |
| **Occluded vessel** |  |
| M1 | 11 [35.5] |
| M2 | 4 [12.9] |
| ACI | 8 [25.8] |
| ACC | 2 [6.5] |
| PCA | 2 [6.5] |
| BAS | 2 [6.5] |
| ACA | 1 [3.2] |
| N.A. | 1 [3.2] |
| **Day** |  |
| d1 ** | 13.0 [8.0-21.0] |
| d2 ** | 14.0 [2.5-31.0] |
| d3 *** | 8.5 [1.5-29.0] |
| d7 # | 7.0 [1.0-33.0] |
| **Thrombus entity** |  |
| Low Shear (TOAST 2) n [%] | 22 [67.7] |
| Undetermined (TOAST 5), n [%] | 6 [19.4] |
| High Shear (TOAST 1, 4) n [%] | 4 [12.9] |
| **Assays performed** |  |
| Flow cytometry based quantitative phenotyping, n | 7 thrombi, 7 blood |
| Flow cytometry based monocyte phenotyping, n | 9 thrombi, 9 blood |
| Flow cytometry based neutrophil phenotyping, n | 8 thrombi, 8 blood |
| FACS-Sort based monocyte PRIME-Seq, n | 7 thrombi, 7 blood |
| FACS-Sort based neutrophil PRIME-Seq, n | 8 thrombi, 9 blood |
| Single-cell RNA Seq, n | 3 thrombi, 3 blood |
| Single-cell CITE Seq, n | 4 thrombi, 3 blood |
| Confocal microscopy, n | 1 thrombus |

\*TOAST 4 thrombi originated from one arterial dissection and one arterial aneurysm with subsequent intra-aneurysm thrombus formation
\*\*n=2 NIHSS scores not available
\*\*\*n=3 NIHSS scores not available
#n=5 NIHSS scores not available

